# Resistance mechanism to Notch inhibition and combination therapy in human T cell acute lymphoblastic leukemia

**DOI:** 10.1101/2022.12.23.521745

**Authors:** Linlin Cao, Gustavo A. Ruiz Buendía, Nadine Fournier, Yuanlong Liu, Florence Armand, Romain Hamelin, Maria Pavlou, Freddy Radtke

## Abstract

Gain-of-function mutations in *NOTCH1* are among the most frequent genetic alterations in T cell acute lymphoblastic leukemia (T-ALL), making the Notch signaling pathway a promising therapeutic target for personalized medicine. Yet, a major limitation for long-term success of targeted therapy is relapse due to tumor heterogeneity or acquired resistance. Thus, we performed a genome-wide CRISPR-Cas9 screen to identify prospective resistance mechanisms to pharmacological NOTCH inhibitors and novel targeted combination therapies to efficiently combat T-ALL. Mutational loss of *Phosphoinositide-3-Kinase regulatory subunit 1 (PIK3R1)* causes resistance to Notch inhibition. *PIK3R1* deficiency leads to increased PIK3/Akt signaling which regulates the cell cycle and spliceosome machinery, both at the transcriptional and post-translational level. Moreover, several therapeutic combinations have been identified, where simultaneous targeting of the cyclin-dependent kinases 4 and 6 (CDK4/6) and NOTCH proved to be the most efficacious in T-ALL xenotransplantation models.

**Key points:**

- Mutational loss of *PIK3R1* induces resistance to NOTCH1 inhibition in T-ALL
- Pharmacological Notch inhibition synergizes with CDK4/6 inhibitors in T-ALL

## Introduction

T cell acute lymphoblastic leukemia (T-ALL) is an aggressive hematological malignancy caused by genetic alterations during T-cell development. The 5-year overall survival rate in pediatric T-ALL patients improved considerably over the past 30 years, whereas it stagnated in adults^1,2^. The increased survival rate is largely due to improved risk-based stratification and applying aggressive combination chemotherapies^3^. However, classical chemotherapy treatment proves inferior when treating relapsed and refractory T-ALL^4^, requiring the implementation of novel therapeutic strategies.

Next generation sequencing (NGS) and related genomic diagnostics of ALL patient samples have not only provided unprecedented insight into different T-ALL subgroups associated with different mutation and gene expression signatures but also identified actionable targets^5,6^. Using this approach, oncogenic gain-of-function mutations in *NOTCH1* have been identified in more than 55% of T-ALL cases^5–7^. The identification of *NOTCH1* as one of the most frequently mutated genes in T-ALL^5–7^ and the finding that *NOTCH* genes are also mutated in other cancers^8^ has boosted the development of a spectrum of therapeutics. These include antibodies neutralizing Notch receptors or ligands, or γ-secretase inhibitors (GSI) preventing NOTCH activation^8^. In this context, we identified and pre-clinically validated a novel orally active small molecule (CB-103) that efficiently blocks the Notch transcription activation complex without causing dose-limiting intestinal toxicities^9^. CB-103 has successfully been evaluated in a recent phase I/II clinical trial (https://clinicaltrials.gov/ct2/show/NCT03422679) and complete response was observed in a patient with relapsed and refractory T-ALL^10^.

Although novel therapeutics blocking Notch signaling show promising outcomes, the use of mono-therapies will likely result in relapse due to tumor heterogeneity and acquired resistance. Thus, a better understanding of resistance mechanisms to Notch inhibitors and the development of combination therapies will facilitate effective treatment of T-ALL patients.

Here, we performed a genome-wide CRISPR-Cas9 screen in human T-ALL cells. We identified the *Phosphoinositide-3-Kinase regulatory subunit 1 (PIK3R1)* as a key player in NOTCH treatment response. Mutational loss of PIK3R1 activity confers resistance to pharmacological Notch inhibition. Unbiased transcriptomic and proteomic analyses in *PIK3R1* deficient T-ALL cells revealed PI3K-AKT mediated up-regulation of pro-survival and proliferation pathways, together with alterations of the spliceosome machinery in response to Notch inhibition. Moreover, our screen led to the identification of pathways that can be pharmacologically targeted synergistically with Notch inhibitors, which resulted in prolonged survival in a preclinical xenograft T-ALL model. Overall, our study identified novel resistance mechanisms to pharmacological Notch inhibition and combination strategies bearing the potential to more efficiently treat refractory or relapsed T-ALL patients.

## Methods

### CRISPR-Cas9 screen and analysis

Cas9-expressing DND-41 cells were transduced with Human GeCKOv2 library (Addgene 1000000048, 1000000049) at MOI~0.3 and selected with 1μg/ml of puromycin for 6 days. Cells were cultured and treated with either DMSO, GSI (DAPT, 10μM) or CB-103 (5μM) for 2 weeks in triplicates. Minimally 3□×□10^7^ cells per replicate were harvested at starting and ending point for genomic DNA isolation. sgRNA sequences were amplified and sequenced using Illumina NextSeq 500. MAGeCK (v.0.5.9.2) was used for data analysis.

### *In vivo* xenografts

All animal experiments were approved by cantonal veterinary service (VD3323, VD3665) and performed in accordance with Swiss national guidelines. Female NSG mice were intravenously injected with 1×10^6^ luciferase-expressing RPMI-8402 cells. Tumor growth was measured twice a week using *in vivo* imaging system (IVIS) and upon tumor establishment, mice were randomized and treated with vehicle or drugs (refer to detailed information in Supplemental data) for two weeks. Tumor growth was monitored until endpoint.

### Proteomics assay

Peptides (250μg) were labeled with Tandem Mass Tags (TMT), pooled and acidified. Phosphopeptides were further enriched using titaium dioxide (TiO_2_) and Ferric nitrilotriacetate. Raw data were processed by Proteome Discoverer (v2.4) against Uniprot Human reference proteome.

### Data sharing statement

Reagents and data will be available upon request to corresponding author.

## Results

### Genome-wide CRISPR screen identifies *PIK3R1* associated with resistance to pharmacological Notch inhibition in T-ALL

The efficacy of targeted therapies for treatment of cancer patients is often limited by development of drug resistance^11^. Potential resistance mechanisms to pharmacological Notch1 inhibition mediated by GSI or CB-103 in T-ALL are currently unclear. Thus, we performed a genome-wide loss-of-function (LoF) CRISPR/Cas9 screen^12^ to identify genes responsible for resistance to Notch inhibition and novel combination therapies for efficient treatment of human T-ALL. We used a Notch-dependent human T-ALL cell line, DND-41, which responds moderately to both GSI and CB-103 treatment *in vitro*^9^. DND-41 cells stably expressing Cas9 were infected with human GeCKO v2 CRIPSR libraries, containing 123,411 sgRNAs targeting 19,050 genes and treated with either vehicle, GSI or CB-103 for 21 days enabling both positive and negative selection of sgRNAs (Figure 1A).

**Figure 1.**
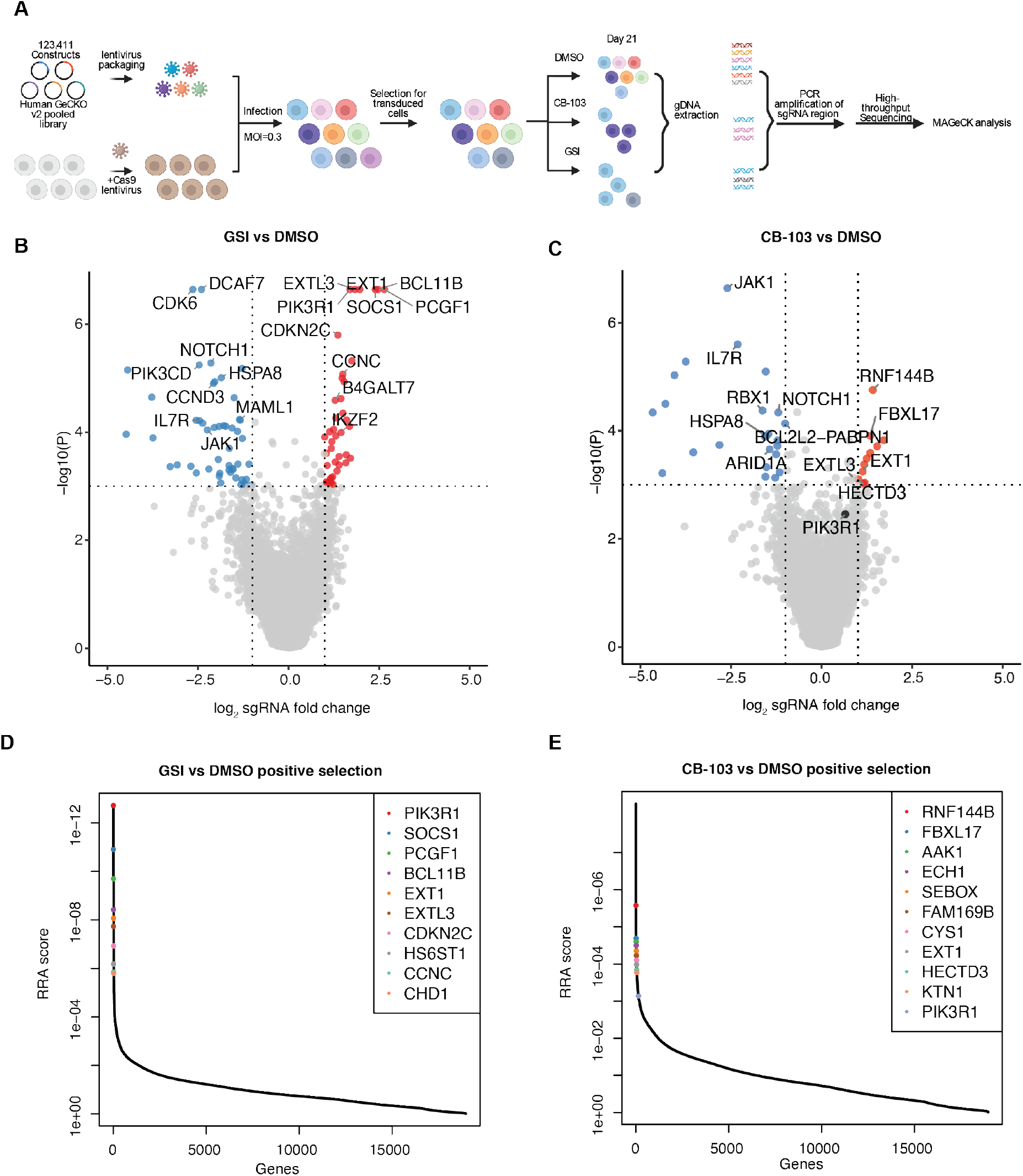
Functional genome-wide CRISPR screen identifies PIK3R1-mediated responsible for resistance to Notch inhibition and druggable candidate pathways for combination therapies in T-ALL. (A) Schematic respresentation of the genome-wide loss-of-function CRISPR screen. (B) Volcano plots depicting gene-targeting sgRNAs negatively or positively selected comparing γ-secretase inhibitor (GSI) vs DMSO treatment. Red, adjusted *P* < 0.001, log_2_ fold-change > 1; blue, adjusted *P* < 0.001, log_2_ fold-change < −1. (C) Volcano plots showing gene targeting sgRNAs negatively or positively selected comparing CB-103 vs DMSO treatment. Red, adjusted *P* < 0.001, log_2_ fold-change > 1; blue, adjusted *P* < 0.001, log_2_ fold-change < −1. (D) Robust rank aggregation (RRA) plots displaying the top 10 enriched sgRNAs comparing GSI vs DMSO treatment. (E) RRA plots displaying top enriched sgRNAs comparing CB-103 vs DMSO treatment.

sgRNAs targeting 293 (GSI-treated) and 131 (CB-103-treated) genes were identified as significantly depleted (P<0.05, log_2_FC<-1) in GSI- and CB-103-treated T-ALL cells compared to vehicle control (Figure 1B-C). Negatively selected sgRNAs indicate genes that, when inhibited, might function synergistically with Notch inhibition to effectively eradicate T-ALL cells. Pathway analysis revealed that significantly depleted genes were regulating MYC- and E2F signaling, as well as G2M checkpoint and mTOR signaling pathways (Supplemental Figure 1).

Conversely, sgRNAs targeting 178 (GSI-treated) and 76 (CB-103) genes were identified as significantly enriched (P<0.05, log_2_FC>1) in GSI- and CB-103-treated cells compared to vehicle control, indicating that the loss of these genes could confer resistance to Notch inhibition. Robust rank aggregation (RRA) method was used to identify genes preferentially lost in response to Notch inhibition (Figure 1D-E). Among these genes, *Phosphoinositide-3-Kinase regulatory subunit 1 (PIK3R1)* was identified at the top of the list in the GSI versus DMSO screen and was also identified in the screen of CB-103 versus DMSO-treated T-ALL cells. The *PIK3R1* gene encodes for the p85α regulatory subunit of PI3K, which contains an SH2 domain that binds to and inhibits the catalytic subunit (p110) of PI3Ks. Interestingly, *PIK3R1* mutations were identified as drivers of tumorigenesis in ovarian cancer^13^, endometrial cancer^14^ and breast cancer^15^. *PIK3R1* hotspot mutations in the SH2 domain were also recently reported in pediatric T-ALL patients^5,6^. In addition, we noticed that the positive regulatory subunit of PI3K (*PIK3CD*, leukocyte-restricted catalytic p110δ subunit) was depleted in Notch inhibitor treated cells. Taken together, these observations suggested a key role of PI3K signaling in acquired resistance to Notch1 inhibition.

### Loss of *PIK3R1* renders T-ALL cells resistant to pharmacological Notch inhibition

To validate the screening results, we generated multiple *PIK3R1* knockout clones in two different NOTCH1-driven T-ALL cell lines (DND-41, RPMI-8402) and stable knockdown clones in the NOTCH3-driven cell line TALL-1 (Supplemental Figure 2 A-C). Loss of *PIK3R1* in several T-ALL cell lines led to a mild growth advantage compared to non-targeting control sgRNA clones (NT) or scrambled shRNA controls (scr). In contrast, cell growth of all GSI- and CB-103-treated *PIK3R1* knock-out (KO) or knock-down (KD) clones was significantly enhanced compared to NT or scr (Figure 2A and Supplemental Figure 2D). We observed a significant decrease in the percentage of cells in S phase in NT or scr T-ALL clones when treated with GSI, confirming that GSI induces cell cycle arrest in T-ALL cells^7^. However, this effect was alleviated in all *PIK3R1* KO and KD cell lines under the same treatment conditions (Figure 2B and Supplemental Figure 2E). Interestingly, we also observed cell cycle arrest in RPMI-8402 and TALL-1 control lines treated with CB-103 and the effect was significantly decreased when *PIK3R1* was lost (Figure 2C and Supplemental Figure 2F). Previously, we showed that CB-103 induces apoptosis in T-ALL cells^9^. Correspondingly, CB-103 treatment for three days induced substantial apoptosis significantly in all three control cell lines. However, loss of *PIK3R1* significantly ablated this effect (Figure 2D and Supplemental Figure 2G). Altogether, these results suggest that loss of *PIK3R1* confers resistance of T-ALL cells to Notch inhibition by protecting them from both drug-induced apoptosis and cell cycle arrest.

**Figure 2.**
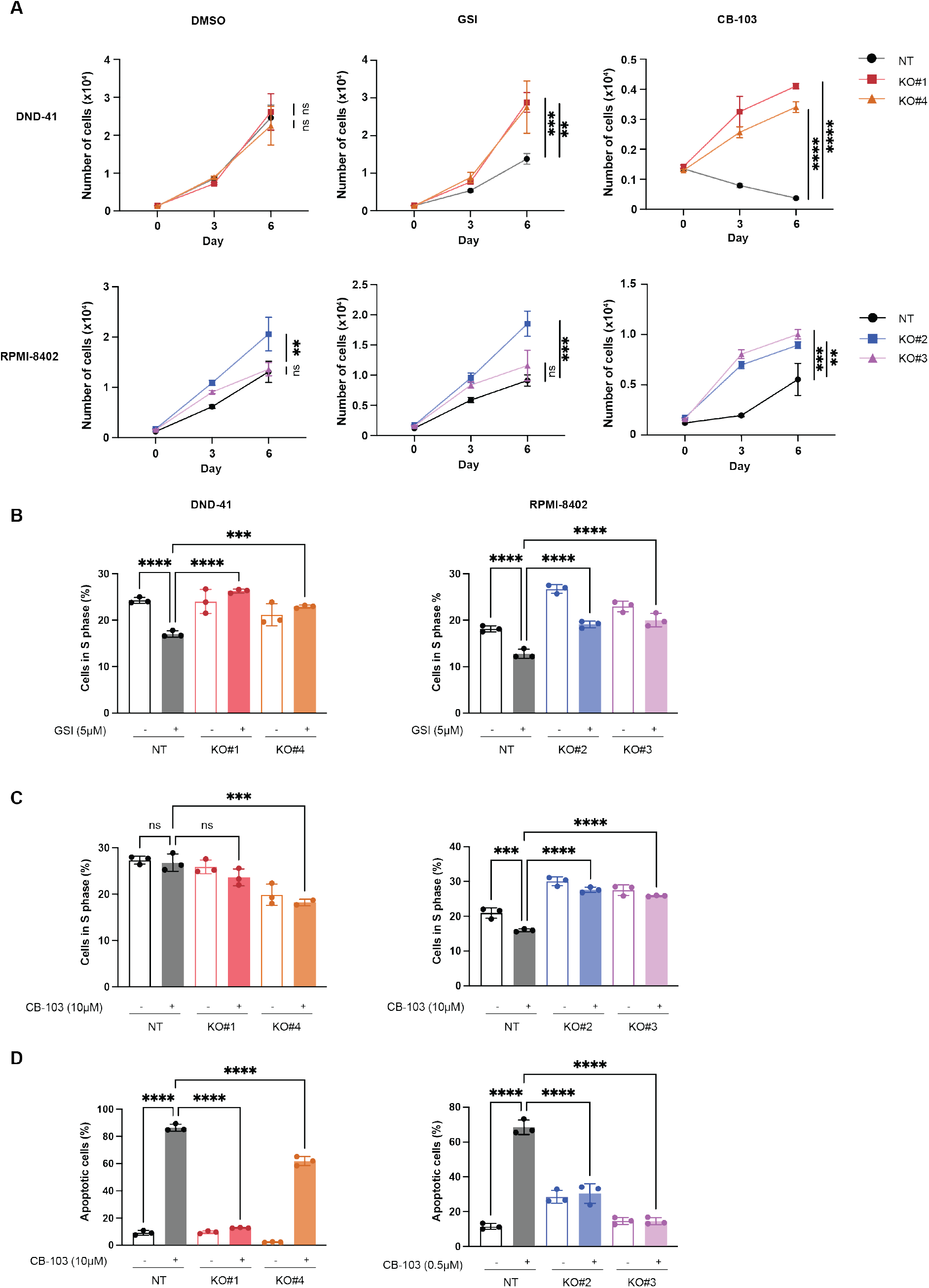
Loss of *PIK3R1* leads to resistance to Notch inhibition in T-ALL cells. (A) Cell proliferation assays of T-ALL *PIK3R1* knockout cell lines under DMSO, CB-103 or GSI treatment conditions. Black connected dots, Non-Target control (NT); colored dots, representative *PIK3R1* knockout cell lines. (B) Cell cycle analyses of *PIK3R1* knockout cell lines performed 6 days post DMSO or GSI treatment at indicated concentrations. (C) Cell cycle analyses of *PIK3R1* knockout cell lines 24hrs post DMSO or CB-103 treatment at indicated concentrations. (D) Apoptosis assays of *PIK3R1* knockout cell lines performed 3 days post DMSO or CB-103 treatment at indicated concentrations. The values shown are mean ± SD (n=3 biologically independent samples, two independent experiments). One-way ANOVA, non-significant (ns), \**P* value < 0.0332, \**\*P* value < 0.0021, \**\*\*P* value < 0.0002, ****P value <0.0001.

### *PIK3R1* deficiency leads to elevated gene expression of proliferation and pro-survival pathways in response to Notch inhibition

To gain insights how loss of *PIK3R1* confers resistance to pharmacological Notch inhibition in T-ALL cells, we performed gene expression analysis (Supplemental Figure 3A). Treatment of RPMI-8402 cells for 24hrs with CB-103 resulted in significantly down regulation of genes associated to hallmark pathways including NOTCH signaling, MYC targets, and E2F targets (Supplemental Figure 3B) as previously reported^9^, whereas GSI treatment resulted in significantly down-regulated MYC targets and MTOR signaling (Supplemental 3B). We did not observe significant enrichment of hallmark pathways analyzing the gene expression differences of RPMI-8402 *PIK3R1* KO versus NT cells, albeit a moderate trend of increased expression of PI3K-AKT and KRAS hallmark pathway genes (Supplemental Figure 3C). This might explain the mild growth advantage observed under normal culture conditions due to loss of *PIK3R1* in T-ALL cells (Figure 2A). Interestingly, Gene Set Enrichment Analysis (GSEA) from CB-103-treated KO versus NT cells revealed an enrichment in multiple hallmarks including E2F targets, MYC targets, PI3K-AKT-MTOR signaling, G2M checkpoint and Apoptosis pathways (Figure 3A). Increased expression of MYC target genes was also observed in GSI-treated KO versus NT cells (Supplemental Figure 3D). Specifically, upregulation of key E2F family transcriptional activators including E2F1, E2F2, E2F3, cell cycle regulators CCND2, CCND3, and down-regulation of the transcriptional repressor E2F5 were observed. In addition, we detected significant up-regulation of anti-apoptotic genes such as BCL2 and BCL-xL (Figure 3B-C). In contrast, typical Notch target genes including *MYC, HES1* or *DTX1* were equally down-regulated in CB-103-treated *PIK3R1* KO and NT cells (Supplemental Figure 3E). These results are consistent with the increased proliferation and survival observed in drug-treated *PIK3R1* KO vs NT cells (Figure 2B-D), and provide some mechanistic insight for Notch inhibitor resistance.

**Figure 3.**
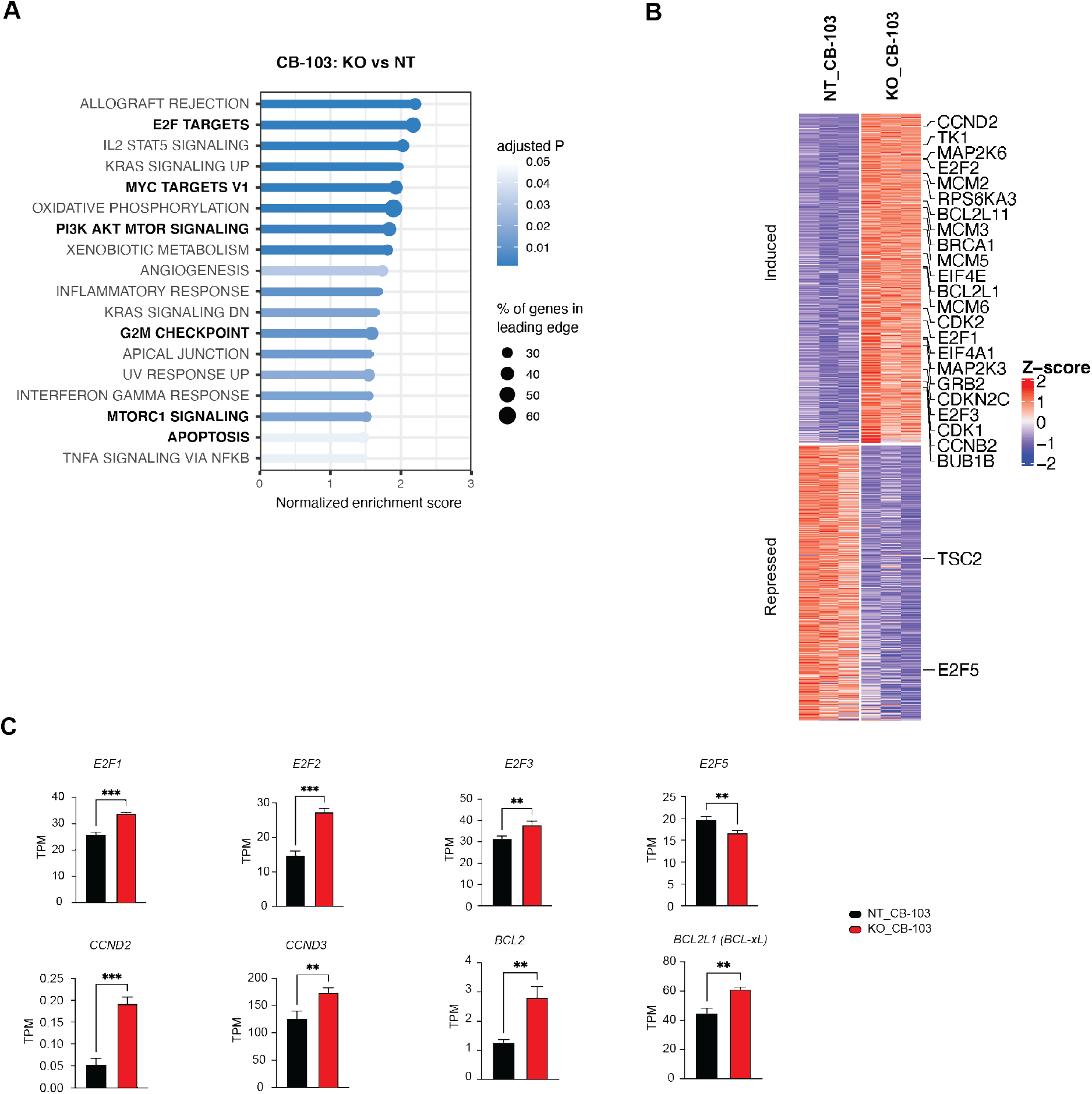
RNA-seq analysis of *PIK3R1* KO cells reveals responses to Notch inhibition at transcriptional level. (A) Top significantly enriched hallmark pathways from a gene set enrichment analysis using differential gene expression results *PIK3R1* KO CB-103-treated compared to NT CB-103-treated T-ALL cells. (B) Heatmap plot showing unbiased clustering of gene expression level changes comparing *PIK3R1* KO CB-103-treated vs NT CB-103-treated T-ALL cells, highlighting key genes involved in E2F targets, PI3K-AKT-mTOR signaling and apoptosis pathway. (C) Expression of a subset of differentially expressed genes measured as transcripts per million (TPM) comparing *PIK3R1* KO CB-103-treated vs NT CB-103-treated T-ALL cells. Values shown are mean ± SD. One way ANOVA test, \**P* value < 0.0332, \**\*P* value < 0.0021, ***P value < 0.0002, ****P value < 0.0001.

### Notch-inhibited *PIK3R1*-mutant T-ALL cells reveal major phosphorylation changes in the cell cycle and spliceosome machinery

The p85 protein encoded by *PIK3R1* is part of an important kinase signaling complex. Its loss may lead to immediate altered signaling events. Therefore, we performed total- and phospho-proteome anaylsis of NT and *PIK3R1* KO cells treated with DMSO or CB-103 (Supplemental Figure 4A and B). Across samples, we quantified 29904 peptides corresponding to 7886 protein groups and 25221 phosphopeptides, of which 21601 were categorized as class I phosphosites^16^ originating from 5531 phosphoproteins (Supplemental Figure 4C). At the total protein level, we observed 54 (NT, CB-103 versus vehicle), 215 (*PIK3R1* KO versus NT) and 206 (*PIK3R1* KO CB-103 versus NT CB-103) significant changes (Supplemental Figure 4D). The comparisons at the phosphorylation level revealed 2983 (NT, CB-103 versus vehicle), 2636 (*PIK3R1* KO versus NT) and 3731 (*PIK3R1* KO CB-103 versus NT CB-103) significant changes (Supplemental Figure 4E). Thus, changes occurring at the level of phosphorylation profiles are much more pronounced compared to changes of the total proteome. KEGG analysis of total protein changes of CB-103-treated PIK3R1 KO versus NT cells, identified cell cycle regulation as the most significantly affected pathway (Supplemental Figure 4F), which corroborated observations from the RNA-seq data. Similar analysis at the phosphoproteome level pointed to cell cycle and spliceosome as the most significant alterations (Figure 4A).

**Figure 4.**
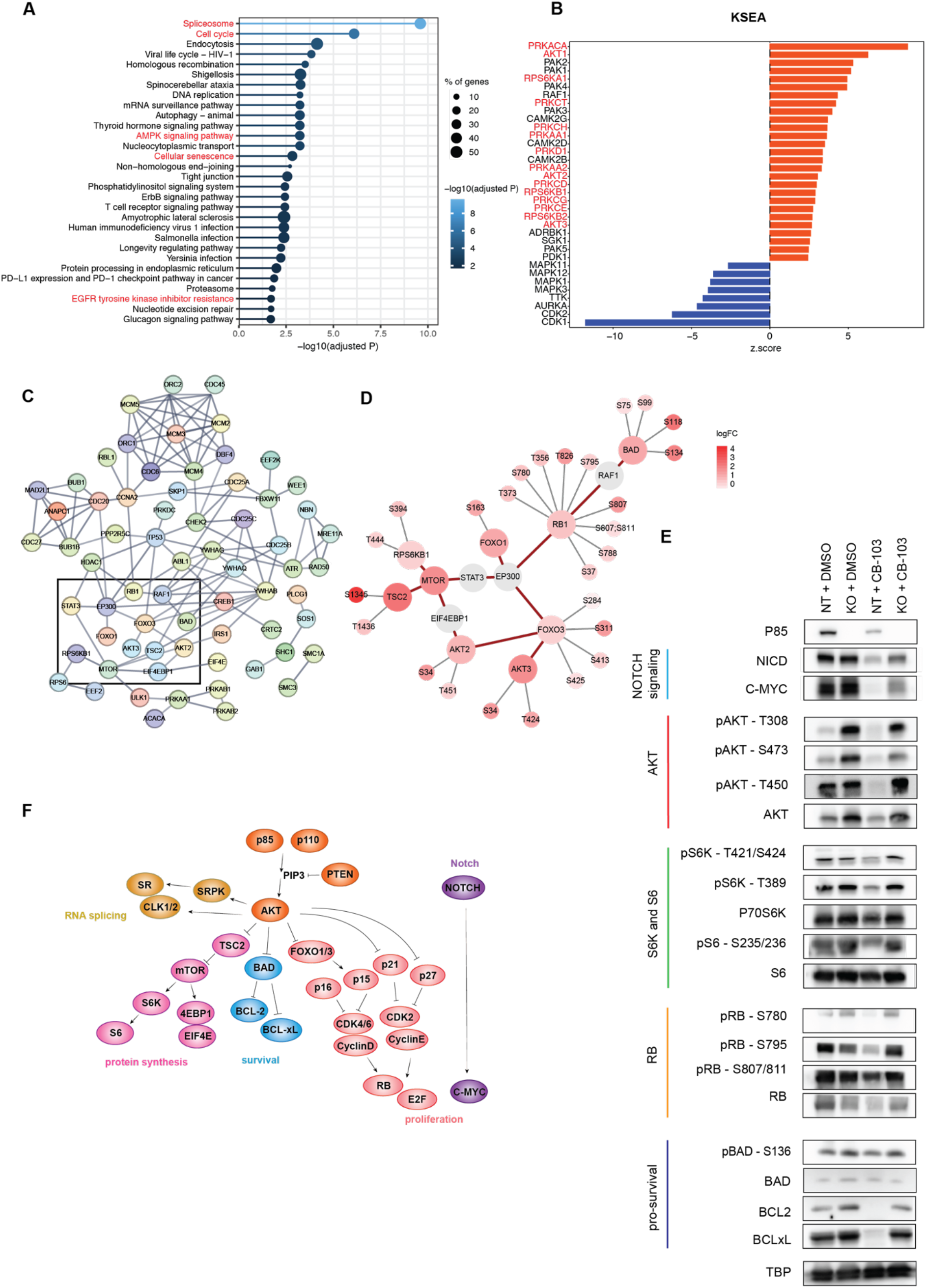
Phosphoproteomics analysis of *PIK3R1* KO cells reveals signaling responses to Notch inhibition. (A) Significantly enriched KEGG pathways of proteins with altered phosphorylation sites comparing *PIK3R1* KO CB-103-treated vs NT CB-103-treated T-ALL cells. Top 30 pathways are shown, solid line color scale indicates adjusted *P* value, dot size of leading edge displays percentage of genes enriched in corresponding pathways. (B) Kinase-Substrate-Enrichment-Analysis (KSEA) of phosphorylation profiles comparing *PIK3R1* KO CB-103-treated vs NT CB-103-treated T-ALL cells. Red, kinases with positive z-score; blue, kinases with negative z-score. (C) Interactions among phosphoproteins within 4 of the top enriched KEGG pathways in (A): assessing cell cycle, AMPK signaling, cellular senescence, EGFR tyrosine kinase inhibitor resistance pathways. Line color indicates the strength of interaction (“Confidence” from the STRING database). Key nodes are gated with black rectangle. (D) Detailed plots of key phosphoproteins with annotated phosphosites and corresponding fold changes from (C) rectangle area. Red circle, identifies phosphoproteins with phosphorylation changes as log_2_ fold-change>1, FDR<0.05. Grey circle, identifies phosphoproteins with phosphorylation sites omitted. Red connecting line, protein interaction from STRING database. Grey radiating line, detailed phosphorylation sites associated with phosphoproteins. (E) Total protein and phosphorylation level of indicated phosphosites by Western blotting for key proteins involved in indicated nodes or pathways: Notch signaling (Light blue); AKT (Red); S6K and S6 (Green); RB (Orange); pro-survival signaling (Dark Blue). TATA-Box binding protein (TBP) was used as loading control. (F) Model summarizing the key nodes of resistance mechanism caused by the loss *of PIK3R1* to Notch signaling inhibition.

To dissect kinase regulation in more detail, we performed Kinase-substrate Enrichment Analysis (KSEA) on the differential phosphorylation profiles of our comparison groups (Figure 4B). The analysis of CB-103 versus vehicle revealed that CB-103 treatment led to decreased AKT1, MTOR, and S6K signaling, whereas the PIK3R1 versus NT comparison showed the expected reciprocal outcome, with increased PKC family, AKT signaling, due to loss of p85, (Supplemental Figure 5A). Importantly, comparison of PIKR1 KO CB-103-treatment versus NT CB-103-treatment showed increased activating phosphorylation events for AKT 1/2/3, PKC family, and S6K, which were maintained and no longer downregulated by CB-103 treatment (Figure 4B).

Subsequently, we examined interactions among key proteins (Figure 4A) using experimentally validated knowledge from the STRING database (Figure 4C) and highlighted phosphorylation changes on these proteins (Figure 4D). This detailed phopho-mapping provides insights regarding functionally established phosphorylation events such as S780 for RB as well as less examined events including T451 on AKT2, which has previously been associated with oncogenic signaling (Figure 4D). Immunoblotting validated key phosphorylation events for AKTs, S6K, RB1 and BAD, which are important regulators of proliferation and cell survival (Figure 4E). CB-103 treatment resulted in marked downregulation of NICD and c-MYC^9^, along with reduced total AKT levels and more pronounced reduced phosphorylation at residues T308, S473 and T450. Yet, these effects were largely ablated in p85-deficient cells (Figure 4E and Supplemental Figure 5B). Similarly, the phosphorylation of ribosome protein S6 kinase (p-S6K, T389, T421/S424) and its downstream substrate S6 (p-S6, S235/236) were down-regulated by CB-103 treatment but not in p85-deficient cells. Thus, loss of *PIK3R1* indeed helps to maintain proteins involved in protein translation under CB-103 treatment. In addition, all phosphorylation sites of RB tested (S780, S795 and S807/811) were down regulated in CB-103 sensitive compared to the resistant cells (RB). The same holds true for BCL2 and BCL-xL, whereas BAD and p-BAD levels (pro-survival) remained comparable. These results confirm that p85-deficient T-ALL cells are able to cope with Notch inhibition through increased AKT signaling and maintain protein translation, cell proliferation and pro-survival pathways.

Interestingly, LoF *PIK3R1* led to prominent phosphorylation changes in proteins involved in the spliceosome and RNA processing in cells treated with pharmacological Notch inhibitors (Figure 4A and Supplemental Figure 6A-B). This analysis allowed to establish changes in phosphorylation profiles of splicing factors upon altered PI3K signaling and highlighted a wide spectrum of so far uncharacterized phosphorylation sites. A recent report linked oncogenic PI3K signaling with splicing alterations in breast cancer on the transcriptional level^17^. Thus we reanalyzed our RNA-seq data for differentially expressed transcripts, which were indeed associated with genes involved in cell cycle and regulation of apoptosis signaling pathways (Supplemental Figure 6C-D). Furthermore, we assessed the differential exon usage using DEXSeq and identified a spectrum of genes with alternative exon usage events in *PIK3R1* deficient cells in response to Notch inhibition compared to NT cells, including transcripts of *ELL Associated Factor 1 (EAF1)* (Supplemental Figure 6E).

Our results show that loss of *PIK3R1* in T-ALL cells led to increased PI3K-AKT signaling, causing major phosphorylation changes in the cell cycle and spliceosome machinery changes that resulted in downstream activation of cell cycle progression, increased cell proliferation, E2F gene activation, increased protein synthesis and cell survival. Changes in the spliceosome at phosphorylation levels correlated also with differential splicing at the transcriptional level. Consequently, these mechanisms contribute to resistance to Notch inhibition in T-ALL (Figure 4F).

### Pharmacological Notch inhibitors synergize with targeted therapies in human T-ALL cells

The advantage using a CRSIPR/Cas9 screen in T-ALL cells under drug selection is that it not only allows for identification of candidate genes mediating drug resistance such as *PIK3R1*, but also genes and pathways crucial for survival under drug selection. This opens avenues to identify novel combination therapies. Preferentially depleted sgRNAs in GSI- and CB-103-treated T-ALL cells pointed to well established signaling components within T-ALL, including components of the IL7/JAK pathway (IL7R, JAK1), regulators of the cell cycle machinery (CDK6:CCND3), and the key gene encoding the PI3K catalytic subunit (*PIK3CD*) (Figure 1B and C).

We validated these candidates using available FDA-approved inhibitors against CDK4/6 (PD-0332991), JAK1/2 (Ruxolitinib), and PIK3δ (CAL-101). We first established *in vitro* sensitivity profiles, and observed that the single agent IC50 of CB-103 for DND-41 cells was 4.3μM and 0.1μM for PD-0332991. We then tested a combination treatment administrating CB-103 and PD-0332991 at three fixed ratios of their corresponding IC50 (1:1, 1:2.5 and 1:0.5) and established dose response curves (Figure 5A). The combination treatment increased sensitivity of the cells to CB-103 by lowering its IC50 to approximately 0.1μM, which is 43-fold lower than single agent treatment (Figure 5A). Similarly, combination of PD-0332991 and GSI lowered the IC50 of GSI approximately 100-fold (Figure 5A). The Combination Index^18^ (CI) was 0.06 for CB-103 plus PD-0332991 and 0.0183 for GSI plus PD-0332991, both of which are below 0.1 indicating very strong synergism (Figure 5B). In addition, combination treatment induces the downregulation of c-MYC which is downstream of Notch and p-RB as key cell cycle regulator in two independent T-ALL cell lines (Figure 5C). Similarly, we observed very strong synergism combining Notch inhibitors with a JAK1/2 inhibitor or a PI3Kδ inhibitor (Figure 5B). These findings suggest that Notch inhibition in combination with FDA-approved compounds targeting CDK4/6, IL7R signaling, or PI3K/AKT pathway should be more efficacious compared to single agent treatment.

**Figure 5.**
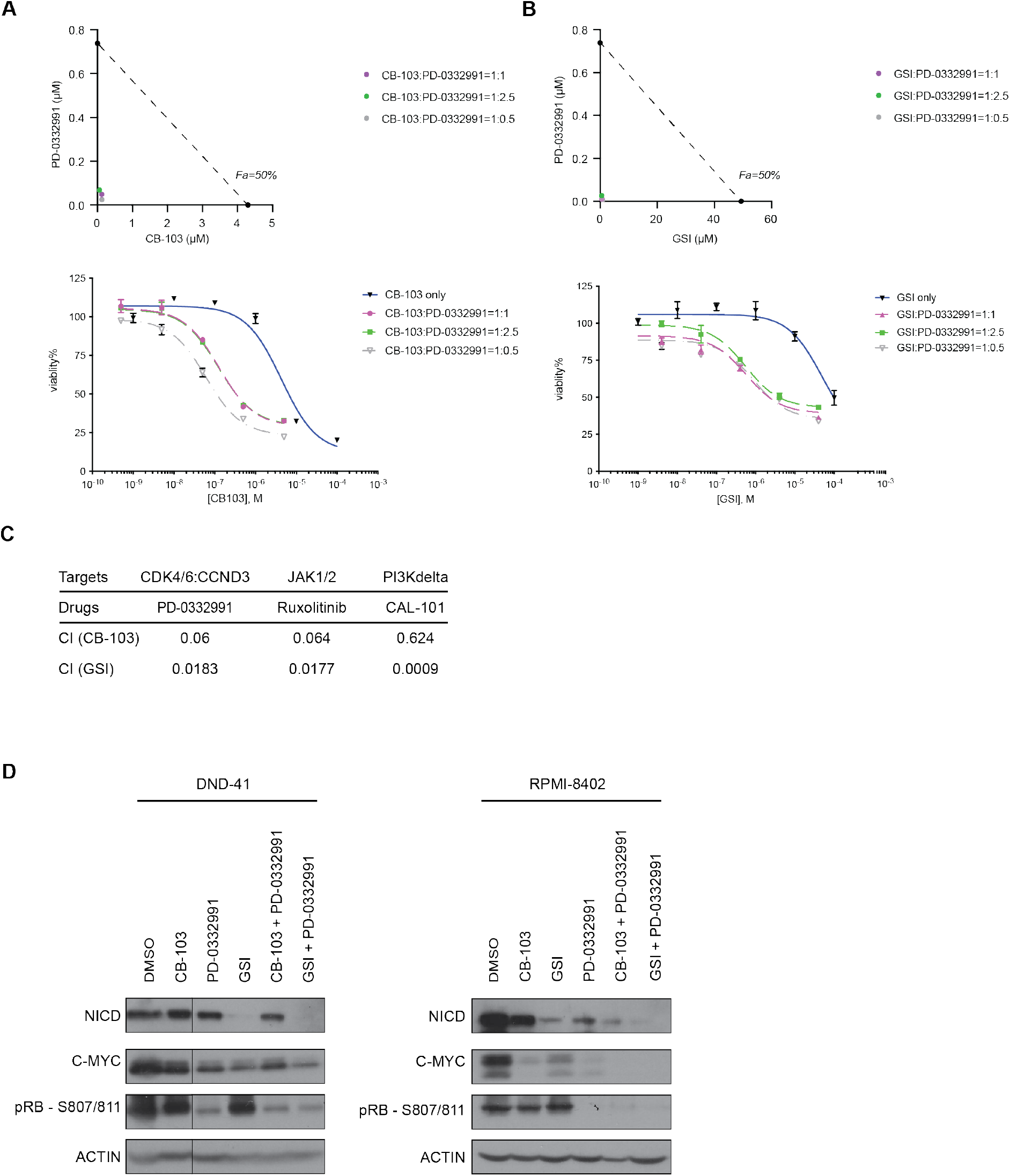
*In vitro* synergy between Notch inhibitors and multiple targeted therapies identified from the CRISPR screen. (A) Isobologram plots (upper panel) and cell survival assay (lower panel) of T-ALL cells in response to CB-103 alone (Blue), or in combination with PD-0332991 with corresponding ratios of each IC50 in decreasing doses for 3 days. Purple, ratio (Notch inhibitor: PD-0332991) = 1:1; Green, ratio = 1:2.5; Grey, ratio = 1:0.5. X-axis plotting concentration of CB-103. The values shown are mean ± SD (n=4 biologically independent samples, two independent experiments performed). (B) Isobologram plots (upper panel) and cell survival assay (lower panel) of T-ALL cells in response to γ-secretase inhibitor (GSI) alone (Blue), or in combination with PD-0332991 with corresponding ratios of each IC50 in decreasing doses for 3 days. Purple, ratio (Notch inhibitor: PD-0332991) = 1:1; Green, ratio = 1:2.5; Grey, ratio = 1:0.5. X-axis plotting concentration of GSI. The values shown are mean ± SD (n=4 biologically independent samples, two independent experiments performed). (C) Table summarizing combination index (CI) of Notch inhibitors together with PD-0332991, Ruxolitinib or CAL-101. (D) Total protein levels of NICD and c-MYC as well as phosphorylation level of p-RB in DND-41 cells (left panel) and RPMI-8402 cells (right panel). Cells were treated with DMSO or corresponding single drugs or drug combinations for 24hrs. ACTIN was used as loading control.

These promising *in vitro* results prompted us to assess their efficacy in xenotransplantation assays. RPMI-8402 T-ALL cells expressing a luciferase reporter were transplanted into NSG mice to monitor tumor growth and progression of disease over time. Animals with established tumors were treated with single agent compounds (vehicle, CB-103, GSI, PD-0332991) or with combination therapy (CB-103 or GSI plus PD-0332991) for two weeks (Figure 6A). The kinetics of tumor progression showed a moderate and statistically significant reduction in tumor burden for both single agent treatments of CB-103 or GSI compared to vehicle (Figure 6B-C). Single agent treatment of PD-0332991 revealed a robust reduction in tumor burden. However, the strongest reduction in tumor burden was observed, when mice were treated with combination of PD-0332991 and either CB-103 or GSI (Figure 6C). To test whether combination treatment led to an increase in overall survival of experimental animals, treatment was ceased after 2 weeks and tumor relapse and survival rates were monitored. Despite the short treatment window, the dual agent treatment of GSI plus PD-033291 translated into significant prolonged overall survival compared to other treatment regiments (Figure 6D).

**Figure 6.**
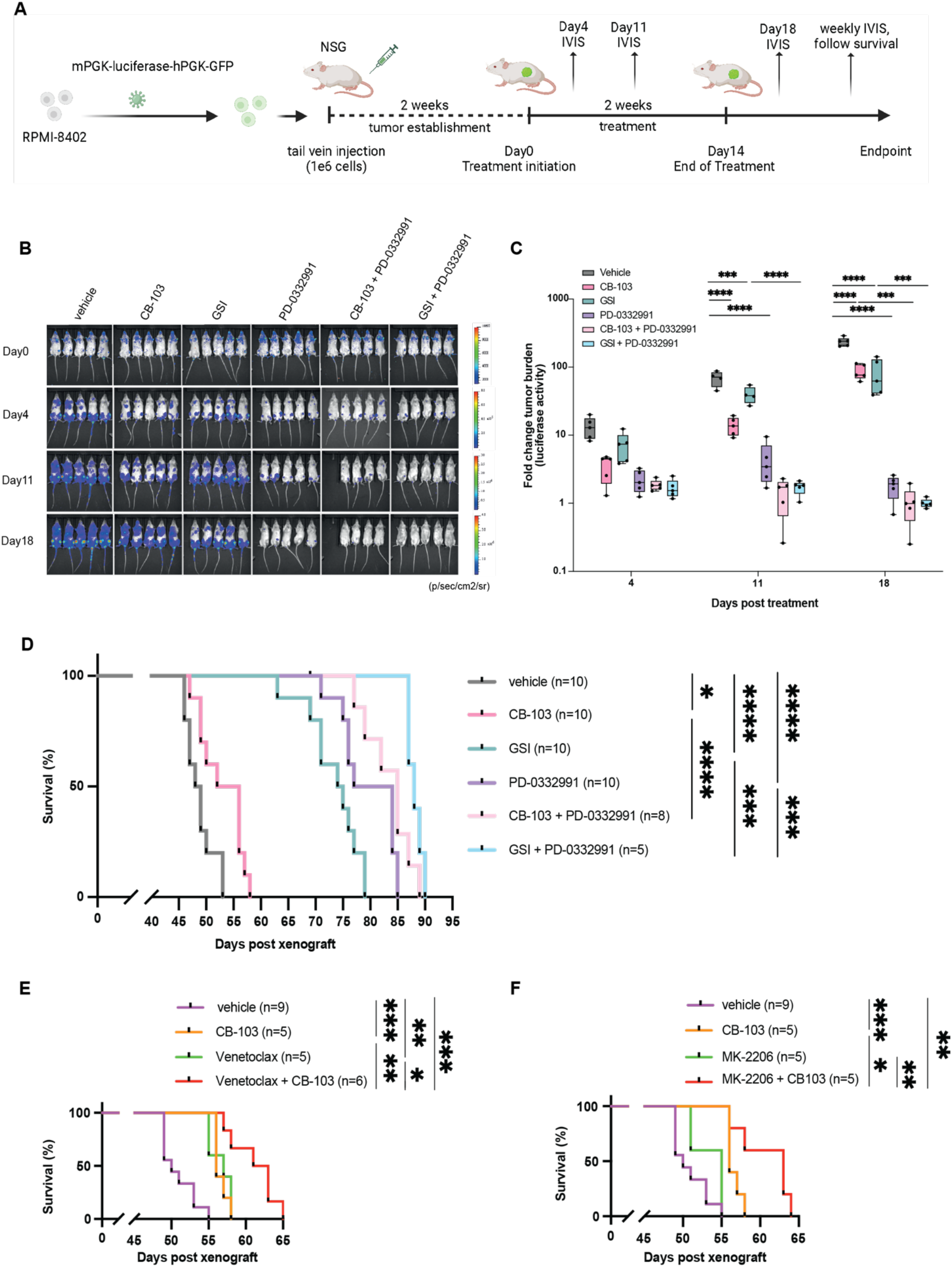
Combination of Notch inhibitors and multiple targeted therapies leads to decreased tumor burden and prolonged survival in human T-ALL cell line xenograft model. (A) Schematic representation of human T-ALL cell line xenograft model and drug treatment study. (B) Representative bioluminescence imaging at days indicated post treatment of each group. (C) Quantification of tumor burden measured by bioluminescent signals at days indicated post treatment of each group, testing Notch inhibitors alone or in combination with PD-0332991. Y-axis shows log_10_ fold change of signals on Day 11 or Day 18 post treatment comparing to initiation of treatment. Data are shown in box and whisker plots showing all data points. One-way ANOVA was performed. (D) Kaplain-Meier survival analysis of NSG mice within each treatment group testing Notch inhibitors, PD-0332991 or in combination. (E) Kaplain-Meier survival analysis of NSG mice within each treatment group, testing CB-103, Venetoclax or in combination. (F) Kaplain-Meier survival analysis of NSG mice within each treatment group, testing CB-103, MK-2206 or in combination. Log-rank (Mantel-Cox) test, *P* value as indicated. \**P* value < 0.0332, \**\*P* value < 0.0021, \**\*\*P* value < 0.0002, \**\*\*\*P* value < 0.0001.

The PI3K-AKT axis was identified as a main switch of downstream signaling events responsible for resistance to Notch inhibition in the CRSIPR/Cas screen, RNA-seq and proteomics data. Therefore, we also tested dual treatment of the AKT inhibitor (MK-2206) combined with CB-103 and observed significant prolongation of overall survival with combination compared to single agent therapy (Figure 6E).

In light of increased BCL2 expression in our RNA-seq data and a recent report on complete clinical response of a relapse refractory T-ALL patient, treated with the BCL2 inhibitor Venetoclax and CB-103^10^, we proceeded to assess the efficacy of combining CB-103 and Venetoclax in our model. Indeed, this combination treatment significantly extended overall survival compared to single agent treatment in a comparable range as with CB-103 plus MK-2206 (Figure 6F). Overall, the CRSIPR/Cas9 screen in T-ALL cells unveiled potentially novel avenues of combination therapies.

## Discussion

NGS analyses of primary T-ALL samples and cell lines has identified the *NOTCH1* as being amongst the most frequently mutated genes throughout different T-ALL subgroups^5,6^. This, together with the identification of gain-of-function mutations in other tumor entities^8^ highlights the Notch pathway as a therapeutic target for precision medicine. However, a major issue with personalized medicine is the establishment of resistance causing relapse. Thus, we performed an unbiased genome-wide LoF CRISPR screen in a NOTCH1-driven T-ALL cell line (DND41). We identified and validated that loss or down-regulation of *PIK3R1* in several human T-ALL cell lines is responsible for resistance to both GSI- and CB-103-mediated Notch inhibition implicating a generic resistance mechanism. Aberrant activation of the PI3K pathway has been demonstrated to contribute to various cancer types including T-ALL^19–22^. Twenty-three to 27% of T-ALL patients harbor mutations in PI3K pathway genes^5,23,24^, raising the question whether all T-ALL patients with activating NOTCH1 mutations and aberrations within the PIK3 signaling cascade might be resistant to Notch inhibitors. A previous report linked PTEN, which negatively regulates PI3K/AKT signaling, in human T-ALL cell lines to GSI-resistance^25^. However, this conclusion is not consistent with observations that GSI sensitivity was comparable in Notch-driven T-ALL cells obtained from wild type and *Pten* deficient mice^26^. Similarly, multiple human T-ALL cell lines carrying mutant *PTEN* alleles are sensitive to GSI^24^. Thus, the loss of PTEN may not a priori be linked to resistance to pharmacological Notch inhibition but might be dependent on the time point amid the T-ALL transformation process. Thus, loss of PIK3R1 or PTEN during drug-mediated selection in a fully established T-ALL may lead to rapid and high activation of the AKT pathway, resulting in continuous proliferation, survival and thus resistance to pharmacological Notch inhibitors, as observed and validated in our genetic CRISPR-based screen. The hypothesis is supported by findings using a mouse model of NOTCH1-induced TALL with subsequent loss of the *Pten* gene once T-ALL has been established. In this model loss of *Pten* indeed resulted in the development of GSI resistance,-unlike the *Pten* knockout models, in which *Pten* was already lost at the onset of Notch mediated disease^26,27^. Therefore, the prediction would be that T-ALL patients with LoF mutations of *PIK3R1, PTEN* or activating mutations in PI3K catalytic subunits at disease onset still respond to Notch inhibition. Nonetheless, individuals that acquire such mutations during treatment or at late-stage disease, are more likely to be resistant to pharmacological Notch inhibitors due to elevated activation of AKT signaling.

To the best of our knowledge, our study provides for the first time a comprehensive analysis of disrupted phosphorylation modification on splicing factors (Supplemental 5B-C) upon altered PI3K signaling in response to Notch inhibition. This was correlated with differential expression levels of transcript isoforms of genes involved in enriched cell cycle and anti-apoptotic pathways as well as with alternative splicing events (Supplemental Figure 6C-E). Recently, oncogenic PI3K signaling was shown to induce expression of alternatively spliced isoforms linked to proliferation and metabolism in breast cancer^17^. The PI3K-AKT pathway has been shown to regulate several proteins of the splicing machinery^28^. Interestingly, genetic alterations in RNA processing factors were identified in 11 % of pediatric T-ALL cases^6^. Inhibitors against SF3B1, a key U2 spliceosome component, which is also differentially phosphorylated (Supplemental Figure 5B-C), have been shown to inhibit growth of T-ALL and other leukemias. Whether treating *PIK3R1* KO T-ALL cells with a SF3B1-inhibitor would re-sensitize them to pharmacological Notch inhibitors or simply kill TALL cells remains to be addressed. Also, the question remains how the altered phosphorylation profile of splicing factors could cause transcript isoform alterations and specifically contribute to the resistance phenotype. One possible explanation is that the subcellular localization and activity of splicing factors might be phosphorylation-dependent and contribute to the expression of particular splice variants involved in key oncogenic pathways. Whether alternative splicing profiles can predict response to Notch inhibition in T-ALL and other cancer contexts requires future exploration.

The complementary part of the study was to identify potential candidate genes or pathways for combination therapies with pharmacological Notch inhibitors. Our CRISPR screen led to the identification of *PIK3CD, IL7R/JAK1*, and *CDK6:CCND3* as potential targets (Figure 1B and C). Unfavored loss of *PI3KCD* (encoding catalytic PI3K subunit P110δ), was notable as its activity is modulated by the negative regulatory subunit p85α, which we identified as a resistance-associated protein to Notch inhibition. Nine percent of T-ALL patients^30^ harbor activating *IL7R* mutations, which causes constitutive JAK1 signaling in T-ALL^31^. Interestingly both *IL-7R* and the downstream mediator *JAK1* were identified to be preferentially lost in the presence of Notch inhibition. Furthermore, we focused on CyclinD3 which has been demonstrated to be essential for T-ALL induction^32^ in a murine model and CDK6, recently identified as molecular target in T-ALL^33^.

We explored a panel of FDA-approved inhibitors and tested them first for their ability to function synergistically with CB-103 or GSI *in vitro*. Interestingly PD-0332991 (inhibiting CDK4/6), Ruxolitinib (inhibiting JAK1/2), and CAL-101 (inhibiting PIK3CD) appear to function synergistically with both CB-103 or GSI (Figure 5B). We further assessed both CB-103 and GSI in combination with PD-0332991, MK-2206 and Venetoclax in xenotransplantation assays. All different combination treatments showed significant prolonged survival when compared to single agent treatment. However, the best combination in terms of overall survival was obtained by combining PD-0332991 with GSI suggesting that simultaneous inhibition of CDK4/6 together with Notch signaling might be worthwhile to be considered in future clinical combination trials.

## Acknowledgements

The authors would like to acknowledge Ute Koch for critical reading, reviewing and editing of the manuscript, the staff from the Flow Cytometry Core Facility (FCCF), Gene Expression Core Facility (GECF) and Center of Phenogenomics (CPG) at EPFL for excellent technical support. We thank Yueyun Zhang for technical help. This work was in part supported by the Swiss National Science Foundation, the Swiss Cancer League.

## Authorship Contributions

**Conceptualization**, F.R.; **Methodology**, LC.; Bioinformatic analysis, G.A.R.B, N.F., Y.L.; **Formal Analysis and Investigation**, L.C.,; Proteomic analysis, F.A., R.H. M.P., **Writing Original Draft**, L.C., F.R.,; **Writing - Review & Editing**, L.C., G.A.R.B., Y.L., M.P., F.R.; **Visualization**, L.C., G.A.R.B., N.F., Y.L; **Supervision**, F.R., **Project Administration**, F.R.; **Funding Acquisition**, F.R.

