## Supplemental material & methods, Tables 1-3, Figures S1-6 for "Resistance mechanism to Notch inhibition and combination therapy in human T cell acute lymphoblastic leukemia" (from another version of this preprint: version 1, 2022-12-23)

**Supplemental Information**

### Supplemental Methods

#### Cell culture and cell lines

All cell lines were maintained in a humidified atmosphere incubator with 5% CO2 at 37°C. The lentiviral packaging cell line HEK293T (RRID: CVCL_0063) was was obtained from D. Trono Lab maintained in DMEM GlutaMAX (Thermo Fisher, Cat#31966047) supplemented with 10% FBS (GIBCO), 10μg/mL Gentamicin (Life Technologies, Cat# 15710049). The human T-ALL cell lines DND-41 and RPMI-8402 were grown in RPMI 1640 GlutaMAX (GIBCO, Cat# 61870010) supplemented with 10% FBS. The human T-ALL cell line TALL-1 was grown in RPMI 1640 GlutaMAX (GIBCO, Cat# 61870010) supplemented with 15% FBS.

#### Lentiviral transduction

For lentiviral transduction of CRISPR vectors, HEK293T cells were seeded at 30-40% confluency in 10cm dishes in DMEM GlutaMAX (Thermo Fisher, Cat#31966047) supplemented with 10% FBS (GIBCO), and transfected the next day with the plasmid of interest and the packaging plasmids VSV-G (RRID: Addgene_8454), and dR8.74 (RRID: Addgene_22036) using Fugene (Promega) or Calcium Phosphate method[1]. Media was changed 16 hours after transfection. Viral supernatant was collected twice, at 48 and 72hours post-transfection, pooled and precipitated overnight at 4°C using Polyethylene Glycol 8000 (PEG-8000) as resin. Then centrifuged at 3000 rpm for 1 hour at 4°C. The pellet was resuspended in cold PBS and used to infect cells in presence of 4μg/μl polybrene (Millipore, Cat#TR-1003-G).

#### Stable cell line generation

*Generation of knockout cell lines by CRISPR/Cas9 using lentiviral transduction.*

For RPMI-8402 *PIK3R1* CRISPR/Cas9 knockout: RPMI-8402 cells were first transduced with lentiCas9-Blast (Addgene, Cat #52962) and selected using blasticidin (Invivogen, Cat #ant-bl-1) to obtain stable a cell line expressing the Cas9 protein. Once stably transduced and selected, cells were transduced with a second lentiviral plasmid containing PIK3R1-targeting sgRNA (1 and 2, please see in Supplemental Table 1, sgRNAs sequences were selected from the top 2 enriched sgRNAs from the screen result), coupled with RFP (Addgene, Cat #57823). Cells were transduced and RFP$^+$

cells were sorted in single cells into 96-well U-bottom plates. Single cells were expanded into clones and screened by Western Blot for loss of p85 expression. KO#1 and KO#2 lines were generated from *PIK3R1*_sgRNA1, KO#3 and KO#4 lines were generated from *PIK3R1*_sgRNA2.

*Generation of knockout cell lines by CRISPR/Cas9 through ribonucleoprotein (RNP) delivery.*

For DND-41 *PIK3R1* CRISPR/Cas9 knockout: DND-41 cells were delivered ribonucleoprotein complexes using IDT protocol (Neon Transfection System). In brief, cells were split one day prior for optimal growth at the time of transfection. Customized Alt-R CRISPR-Cas9 crRNAs and Alt-R CRISPR-Cas9 tracrRNA ATTO550$^+$ (IDT, Cat #1075927) were annealed to duplex, and then introduced to cells together with Alt-R HiFi Cas9 Nuclease V3 (IDT, Cat #1081060) in Resuspension Buffer R (Neon System Kit). ATTO550$^+$ cells were sorted 24hrs post-electroporation in single cells into 96-well plates. Single cells were expanded to colonies and clones were selected as described above. KO#1 and KO#2 lines were generated from *PIK3R1*_crRNA1, KO#3 and KO#4 lines were generated from *PIK3R1*_crRNA2. NT lines were generated from Alt-R® CRISPR-Cas9 Control Kit (Cat #1072554), Human with same procedure. (Oligo sequences are listed in Supplemental Table 1).

*Generation of stable knockdown cell lines through shRNA.*

For TALL-1 *PIK3R1* knockdown: TALL-1 cells were transduced with hU6_mir30-shRNA-hPGK_GFP-P2A-Luc2-P2A-BlaR (gift from Dr. J. Huelsken Lab), GFP$^+$ cells were sorted in bulk and cell lines were screened by Western Blot for loss of p85 expression. The KD#5 line was generated from *PIK3R1*_shRNA5, KD#8 was generated from *PIK3R1*_shRNA3/4/5/6 in combination. shRNAs sequences were chosen based on previous published data[2] and Sigma Mission Human shRNA website (oligo sequences are listed in Supplemental Table 1).

**Proliferation assay (using flow cytometry)**

Cells were seeded at same numbers in appropiate media (vehicle or drug treated) and cultured in 96-well U-shaped plates for 6 days, in 3-4 replicates. At indicated time points, live cell numbers were counted counted using Attune NxT (ThermoFisher) and data was analyzed using FlowJo v.10.8.1.

**Cell cycle analysis**

Cells were permeabilized with 0.01% Triton-X100 on ice for 3hours, fixed and stained with FxCycle PI/Rnase staining solution (Invitrogen, Cat #F10797) overnight at 4$^{o}$C. Data acquisition was performed on Gallios (Beckman Coulter) and analyzed using FlowJo v.10.8.1.

**Apoptosis analysis**

Cells were stained with the anti-Cy5-Annexin V (1:100, BD Biosciences, Cat#559934, RRID: AB_2869267) and 7-aminoactinomycin D (7-AAD) (1:20, BD Bioscience, Cat #559925, RRID: AB_2869266) in AnnexinV binding Buffer (Biolegend, Cat #422201). Data acquisition was performed on Gallios (Beckman Coulter) and analyzed using FlowJo v.10.8.1.

**Immunoblotting**

Cells were washed twice with 1x PBS and resuspended in RIPA buffer (50mM Tris-HCl, pH 7.5, 150mM NaCl, 1% Nonidet P-40, 0.5% sodium deoxycholate and 0.1% SDS) supplemented with PhosSTOP (Sigma Aldrich, Cat #4906837001) and Protease inhibitor (Sigma Aldrich, Cat #P8340). Whole cell lysates were separated on 10% Tris-Glycine (SDS) gel and transferred to polyvinylidene difluoride (PVDF; Sigma Aldrich, Cat #3010040001) membranes. Membranes were incubated with the indicated primary antibodies overnight at 4$^{o}$C (Supplemental Table 2). Washed membranes were incubated with secondary antibody for 1hour at room temperature. Membranes were developed using Pierce ECL Western Blotting Substrate (Thermo Fisher Scientific, Cat# 32209).

***In vitro* synergy analyses.**

A total of 5,000 cells per well were seeded in a final volume of 100μl with the indicated amount of each drug either alone or in combination in 96-well flat bottom plates. Each treatment was carried out in 3-4 replicates for 3 days. 10μl of AlamarBlue (Life Technologies, Cat #DAL1025) was added to each well and incubated for 4 hours followed by plate reading using Infinite F500 multimode reader (Tecan). Values were normalized to those of no treatment controls and analyzed in GraphPad Prism 7.0

applying non-linear regression dose-response curve calculations to obtain the IC50 values. Combination index (CI) was calculated according to the Chou-Talalay algorithm[2] using the formula: CI = [D]1/[Dx]1 + [D]2/[Dx]2. [D]1 and [D]2 are the concentrations of drug 1 and drug 2 to show a certain effect when treated with two drugs together. [Dx]1 and [Dx]2 are the concentrations that show the same effect with a combination of drug 1 and drug 2 when treated with each drug alone. Synergism can be defined as follows: CI < 1 indicates a synergistic effect; CI = 1 indicates an additive effect; CI > 1 indicates an antagonistic effect.

***Animals and in vivo xenotransplantation***

NSG (NOD.*Cg-Prkdz$^{scid}$ IL2rg$^{tm1Wjl}$*/Szj; SN 005557) mice were originally purchased from Jackson laboratory. RPMI-8402 cells were engineered to express luciferase by lentiviral transduction using mPGK-Luciferase-hPGK-eGFP plasmid. One million cells were intravenously injected through the tail vein into female NSG mice. Tumor growth was measured twice a week using an *in vivo* imaging system (IVIS, Caliper Life Sciences). Briefly, 12 min after intraperitoneal administration of 150mg/Kg of D-luciferin (Biosynth, Cat #L-8220), mice were anesthetized in an induction chamber ($O_2$ and 2% isoflurane) and transferred to the imager. Images were acquired and analyzed using Living Image Software v4.4 (Caliper Life Sciences). Upon tumor establishment (~14 days post xenograft transplantation), animals were randomized based on tumor burden into either vehicle or drug treatment groups. Tumor growth was continuously monitored at indicated time points until endpoint.

CB-103, dissolved in castor oil by sonication, was administrated intraperitoneally once per day (40mg/Kg). LY3039478, dissolved in solvent (93% water, 5% ethanol and 2% Tween80) and administrated intraperitoneally once per day (20mg/Kg). PD-0332991, dissolved in 50mM sodium lactate at pH4, was administrated by oral gavage once per day (150mg/Kg). MK-2206, dissolved in 30% Captisol, was administrated by oral gavage twice a week (120mg/Kg). Venetoclax, dissolved in 35% PEG400, 0.5% Tween80 and 64.5% water, was administrated by oral gavage once per day (35mg/Kg). Drugs were administrated for 14 days and mice were monitored for tumor development according to scoresheet. (For detailed vendor information please see in Table 3.)

**Ethics statement**

All animal work was carried out in accordance with Swiss national guidelines. This study (VD3323, VD3665) was reviewed and approved by the cantonal veterinary service (Service vétérinaire cantonal de Vaud).

**CRISPR-Cas9 screen and analysis**

The Human GeCKOv2 CRISPR knockout libraries A and B (Addgene #1000000048, #1000000049) were pooled and used to identify genes responsible for Notch resistance in T-ALL cells. In brief, a stable Cas9-expressing T-ALL cell line (DND-41-Cas9) was established by lentiviral transduction of DND-41 cells with LentiCas9-Blast (Addgene, Cat #52962) and selected using 10ug/mL Blasticidin (Invivogen, Cat #ant-bl-1). Stable expression of Cas9 protein in the DND-41-Cas9 T-ALL cell line was verified by Western Blot analysis. Subsequently, these cells were transduced with GeCKO v2A library containing 123,411 unique sgRNA sequences targeting 19,050 human genes at a low MOI (~0.3) to ensure effective barcoding of individual cells. Next, transduced cells were selected with 1µg/mL of puromycin (Life Sciences, Cat #A1113803) for 6 days to generate a mutant cell pool. A cell aliquot was frozen as Day0 sample. Cells were cultured and treated with either vehicle (DMSO), $\gamma$-secretase inhibitor (GSI) (N-[N-(3,5-Difluorophenacetyl)-L-alanyl]-S-phenylglycine t-butyl ester) (DAPT, 10$\mu$M) or CB-103 (5$\mu$M) for 2 weeks for sgRNA selection with each condition done in triplicates. At endpoint, at least 3x$10^7$ cells per replicate were collected for genomic DNA extraction to ensure over 400-fold coverage of library using ZymoResearch Quick-gDNA MidiPrep Plus Kit (Zymo Research, Cat#ZYM-D4075-25TST). sgRNA sequences were amplified in two rounds: first round by GeCKO-F1/R1 primers using Herculase II Fusion DNA polymerase (Agilent Technologies, Cat #600679); second round by NGS-Lib_Fwd/Rev 1-10 indexed using NEBNext® High-Fidelity 2X PCR Master Mix (Cat #M0544L). PCR products were purified (Qiagen gel purification kit, Cat #28706, and Qiagen PCR purification kit, Cat #28106), quantified by Qubit (Thermo Fisher Scientific), and profiled by Fragment Analyzer.

Samples were sequenced in one run on Illumina NextSeq 500 Instrument (Illumina) according to the manufacturer instructions, yeilding 20 to 48 mio single-end 75 nucleotide reads per sample. MAGeCK[3] (v.0.5.9.2) was used to analyse the data. Reads were trimmed of their adapters with bcl2fastq v2.19 (Illumina) and quality-controlled with fastQC v0.11.5. Read count tables were obtained using the mageck count command with default parameters on the Human GeCKOv2 combined library of A and B[3]. Treatments were compared to untreated DMSO/vehicle samples using the mageck test command with default parameters and pooling all replicates. Robust Rank Aggretion (RRA) tables were generated. Genes with an adjusted $P < 0.001$ and a $\log_2$ fold-change > 1 or < -1 were considered as positive and negative significant hits, respectively.

Gene set enrichment analysis (GSEA)[4] was performed by ranking genes according to their RRA values in descending order. GSEA was performed with the clusterProfiler R package (v4.0.5)[5] using 10,000 permutations, minimum gene set size = 10, maximum gene set size = 1,000, and hallmark gene sets[6] from the msigdbr package (7.4.1)[7]. Pathways with an adjusted $P < 0.05$ were deemed as significantly enriched. If more than 20 pathways were significantly enriched, the top 20 are shown in the corresponding dotplots.

**RNA-seq sample preparation and data analysis**

RNA was extracted from cell lines using RNeasy Mini Kit (QIAGEN, Cat#74104). RNA quality was assessed on a Fragment Analyzer (Agilent) or TapeStation 4200 (Agilent) to proceed to the polyA-based stranded mRNA library prep method (scores >7.2). Libraries for mRNA-seq were prepared with the Stranded mRNA Ligation method (Illumina) starting from 1µg RNA, according to manufacturer's instructions. Libraries, all bearing unique dual indices, were subsequently loaded at 250pM in a HiSeq 4000 instrument (Illumina) and sequenced according to manufacturer's instructions, yielding paired-end reads of 75 nucleotides. Reads were trimmed of their adapters with bcl2fastq v2.20 (Illumina) and quality-controlled with FastQC v0.11.9.

Reads were aligned to the human genome build hg38 using HISAT2 (v2.2.1)[8]. Transcript-level counts and transcripts per million (tpm) were estimated using Salmon (v1.4.0). Counts were then summarized at the gene level with the tximport R package (v1.20.0)[9]. Genes with an average expression per condition less than counts per million reads mapped (cpm) = 1, and a coefficient of variation per condition higher than 100% were removed from all conditions with the filtered.data() function from the NOIseq R package (v2.36.0)[10] using the following parameters: cv.cutoff = 100, cpm

= 1. Reported $log_2$ fold-change values are the average of the $log_2$ fold-changes computed by edgeR (v.3.34.0)[11,12], DESeq2 (v.1.32.0)[13] and voom from the limma R package (v.3.48.1)[14]. Three differential gene expression analysis methods were run in parallel with the consensusDE R package (v1.14.0)[15]. Differentially expressed genes were defined as genes with Benjamini-Hochberg-adjusted $P < 0.05$. Genes were ranked according to their $log_2$ fold-change in descending order and GSEA was performed with the GSEA() function from the clusterProfiler R package with the following parameters: minGSSize = 10, maxGSSize = 1000, eps = 0, pvalueCutoff = 0.05. Pathways with an adjusted P < 0.05 were deemed as significantly enriched. If more than 20 pathways were significantly enriched, the top 20 are shown in the corresponding dotplots.

*Differential transcript expression*

Transcript expression was quantified with Salmon (v.1.4.0). Transcript-level quantification estimates were imported to R with the tximport package (v.1.22.0). Low-expression transcripts with less than 10 reads were filtered from dataset. Subsequent differential transcript expression analysis was performed with DESeq2 (v.1.34.0) and significantly differentially expressed transcripts were considered as those with an adjusted $P < 0.01$ and an absolute $log_2$ fold-change > 1. Pathway enrichment analysis was performed by selecting the top 100 most significant up- and downregulated transcripts, obtaining the genes they correspond to and using that unique set of genes in an over-representation analysis (ORA) with the enricher() function from clusterProfiler (v.4.2.2) and gene sets obtained with the msigdbr package (v.7.5.1). Pathways with an adjusted $P < 0.05$ were considered as significantly enriched. If more than 20 pathways were significantly enriched, the top 20 are shown in the corresponding plots.

*Differential exon usage*

Differential exon usage analysis using RNA-seq data was performed with the DEXSeq package (v.1.40.0). For each sample, BAM files were converted to SAM format with samtools (v.1.9). The python script available with DEXSeq called "dexseq_prepare_annotation.py" was used to prepare a flattened annotation file in GFF format by using the same GTF file used in the alignment step. Next, the number of reads overlapping each exon counting bin defined in the GFF file was obtained with the python script provided with DEXSeq called “dexseq_count.py” using the following parameters: -p yes, -r pos to indicate respectively that the data was from a paired-end sequencing run, and that the alignment data was sorted by position. For each comparison, the corresponding count files were used to proceed with the downstream analysis steps where a DEXSeqDataSet object was constructed and used for the normalization, dispersion estimation, and differential exon usage steps with default parameters. The parameter “design = ~ sample + exon + condition:exon” was used each time to define the full model formula. Significant differentially used exons were considered as those with a false discovery rate of 10%. Exon usage plots from significantly differentially used exons were obtained with the DEXSeqHTML() function.

**Sample preparation for proteomics study**

Proteolytic digestion was performed with Lys-C (FUJIFILM Wako Pure Chemical Corporation) in an enzyme/protein ratio of 1:100 (w/w) for 4 hours followed by a 6-fold dilution with 50mM $(NH_4)HCO_3$ pH 8 (Sigma) to 1M GndCl. Samples were further digested with Trypsin gold (Promega) 1:40 overnight at 37°C. Resulting peptides were desalted using a 100mg SEP-PAK C18 cartridge (Waters) and vacuum centrifuged. For Tandem Mass Tag (TMT) labeling, dried peptides from each sample (250µg) were first reconstituted in 30µl 100mM HEPES pH 8 and 12µl of TMT solution (66,6µg/µl in pure acetonitrile) was then added. TMT labeling was performed at room temperature for 1.5h and reactions were quenched with hydroxylamine to a final concentration of 0.4% (v/v) for 15min. TMT-labeled samples were then pooled at a 1:1 ratio across all samples. A single shot control LC-MS run was performed to ensure similar peptide mixing across each TMT channel to avoid the need of further excessive normalization. The combined samples were then desalted using a 500mg SEP-PAK C18 cartridge (Waters) and vacuum centrifuged. For proteome analysis of TMT-labeled samples, 5% of the pooled samples were fractionated into 12 fractions using an Agilent OFF-Gel 3100 system following the manufacturer’s instructions. Resulting fractions were desalted on SDBRPS StageTips[16] and dried by vacuum centrifugation. Phosphopeptides were enriched using the sequential metal oxide affinity chromatography (SMOAC) strategy using two consecutive High Select TiO2 enrichments and a Fe-NTA enrichment (Thermo Scientific). Eluates were immediately acidified, combined and dried by vacuum centrifugation. Resulting phosphopeptides were fractionated with the Pierce High pH Reversed-Phase Peptide Fractionation Kit (Thermo Scientific). All fractions were dried by vacuum centrifugation and stored at −20°C until ~~the~~ day of mass spectrometry assessment and analysis.

**Mass spectrometry analysis**

Mass spectrometry assay and analysis were performed to standard protocol (Ref here). Shortly, each individual fraction was resuspended in 2% acetonitrile; 0.1% formic acid and nano-flow separations were performed on a Dionex Ultimate 3000 RSLC nano UPLC system on-line connected with a Lumos Fusion Orbitrap Mass Spectrometer interfaced with FAIMS Pro. A capillary precolumn (Acclaim Pepmap C18 ; 3µm-100Å ; 2cm x 75µm ID) was used for sample trapping and cleaning. Analytical separations were performed at 250nl/min over a 150-min biphasic gradient on a 50cm long in-house packed capillary column (75µm ID; ReproSil-Pur C18-AQ 1.9µm silica beads; Dr. Maisch). Acquisitions were performed through Top Speed Data-Dependent acquisition mode using 3 seconds cycle time. First MS scans were acquired at a resolution of 120’000 (at 200m/z) and the most intense parent ions were selected and fragmented by High energy Collision Dissociation (HCD) with a Normalized Collision Energy (NCE) of 37.5% using an isolation window of 0.7m/z. Fragmented ions scans were acquired with a resolution of 50’000 (at 200m/z) and selected ions were then excluded for the following 120s.

**Proteomics data analysis**

Raw data were processed using SEQUEST, Mascot, MS Amanda[17] and MS Fragger[18] in Proteome Discoverer v.2.4 against the Uniprot Human reference proteome (77027 canonical and isoform Sequences - Last Modified 21/01/29). Enzyme specificity was set to Trypsin and a minimum of six amino acids was required for peptide identification. Precursor tolerance and ion fragment tolerance were set at 10 ppm and 0.02 Da, respectively. Up to two missed cleavages were allowed and a 1% FDR cut-off was applied both at peptide and protein identification levels. For the database search, carbamidomethylation (C) and TMT tags (K and Peptide N termini) were set as fixed modifications whereas oxidation (M) and phosphorylation (S,T,Y) were considered as a variable. Proteome Discoverer node ptmRS was used for analysis and mapping of peptide/protein phosphorylation sites.

The resulting data were processed through in-house written R scripts (v.4.1.2)[19]. The Proteome Discoverer proteins and peptide tables were processed separately. Common contaminants and proteins with a low FDR confidence were filtered out so that the proteins were not quantified in all the TMT channels. The TMT abundances were $log_2$ transformed.

*Normalisation and Limma analysis of the proteome*

Proteins with less than 2 peptides were discarded. A first normalization step was applied according to the Sample Loading normalization[20]. Assuming that the total protein abundances were equal across the TMT channels, the reporter ion intensities of all spectra were summed and each channel was scaled according to this sum, so that the sum of reporter ion signals per channel equals the average of the signals across samples. Following this, a trimmed M-mean (TMM) normalization step was also applied using the package EdgeR[11] (v.3.34.1). The normalization factors obtained from each step were used for the normalization of the phosphopeptide sets. Differential protein expression analysis was performed using R bioconductor package limma (v.3.50.0)[14], followed by Benjamini-Hochberg multiple-testing method[21].

*Normalisation and limma analysis of the phosphosites*

The peptides with phosphosite(s) assigned with a confident localization probability greater or equal to 0.75 were retained. The intensities of phosphopeptides carrying different PTMs (apart from the common phosphorylation) were summed. The TMT abundances of the phosphopeptides were then normalized using the normalization factors originating from the proteome analysis. Differential protein expression analysis was performed using R bioconductor package limma (v.3.50.0)[14], followed by Benjamini-Hochberg multiple-testing method[21].

*Definition of differentially expressed proteins and differentially phosphorylated sites*

A protein was defined as differentially expressed with a threshold of adjusted $P < 0.05$ and abs ($log_2$ fold-change) > 0.5. Differentially phosphorylated sites were defined with a threshold of 1) being situated on Class I phosphopeptides, 2) adjusted $P < 0.05$, and abs ($log_2$ fold-change) > 1, and 3) either its absolute differentially expression level is higher than that at the corresponding proteome level or its fold-change is on the opposite direction than total proteome change.

*Phosphosite visualization*

Protein-protein interactions were retrieved from the STRING database (v11.5)[22]. Only those interactions having active interaction sources from "Experiments" were kept. Minimum required interaction score is 0.700. Network nodes were colored by the average $log_2$ fold-change of displayed phosphosites.

*Pathway enrichment analysis*

We performed pathway enrichment analysis for differentially expressed proteins and for differentially phosphorylated proteins. We used the clusterProfiler R package (4.0.5)[5] on Gene Ontology (GO) and KEGG terms separately. Pathways with an adjusted $P < 0.05$ were deemed as significantly enriched.

**Software**

Quantitative data and statistical analysis were performed with GraphPad Prism v.9.4.1 (RRID: SCR_002798). Flow cytometric data were analyzed with FlowJo v10.8.1 (RRID: SCR_008520). Representation of data was generated in Adobe Illustrator 2021 (RRID: SCR_010279).

### Supplemental Tables
### Supplemental Table 1 – Oligos

| Name | sequence (5'->3') |
|---|---|
| PIK3R1 sgRNA_1F | CAC CGA CTG AGC TAG AGA TTC ATT C |
| PIK3R1 sgRNA_1R | AAACGAATGAATCTCTAGCTCAGTC |
| PIK3R1 sgRNA_2F | CACCGCGCTTTCAAACGCTATCTCC |
| PIK3R1 sgRNA_2R | AAACGGAGATAGCGTTTGAAAGCGC |
| NT_sgRNA_1F | CACCGCTGAAAAAGGAAGGAGTTGA |
| NT_sgRNA_1R | AAACTCAACTCCTTCCTTTTTCAGC |
| PIK3R1 crRNA_1 | rArCrUrGrArGrCrUrArGrArGrArUrUrCrArUrUrCrGrUrUrUrUrArGrArGrCrUrArUrGrCrU |
| PIK3R1 crRNA_2 | rCrGrCrUrUrUrCrArArArCrGrCrUrArUrCrUrCrCrGrUrUrUrUrArGrArGrCrUrArUrGrCrU |
| PIK3R1-sh3F | AGCGATTCAACCACAGAACTGAAGGAATAGTGAAGCCACAGATGTATTCCTTCAGTTCTGTGGTTGAAT |
| PIK3R1-sh3R | GGCAATTCAACCACAGAACTGAAGGAATACATCTGTGGCTTCACTATTCCTTCAGTTCTGTGGTTGAAT |
| PIK3R1_sh4F | AGCGATGTCTTCTCATGATGGGAATAGTGAAGCCACAGATGTATTCCCATCATGAGAAGACAT |
| PIK3R1_sh4R | GGCAATGTCTTCTCATGATGGGAATACATCTGTGGCTTCACTATTCCCATCATGAGAAGACAT |
| PIK3R1_sh5F | AGCGGCGGTACAGCAAAGAATACATAATAGTGAAGCCACAGATGTATTATGTATTCTTTGCTGTACCGC |
| PIK3R1_sh5R | GGCAGCGGTACAGCAAAGAATACATAATACATCTGTGGCTTCACTATTATGTATTCTTTGCTGTACCGC |
| PIK3R1_sh6F | AGCGCCGAGCCCTATAACTTGTACAAATAGTGAAGCCACAGATGTATTTGTACAAGTTATAGGGCTCGG |
| PIK3R1_sh6R | GGCACCGAGCCCTATAACTTGTACAAATACATCTGTGGCTTCACTATTTGTACAAGTTATAGGGCTCGG |
| scr_F | AGCGTAGCGACTAAACACATCAAAATAGTGAAGCCACAGATGTATTTTGATGTGTTTAGTCGCTA |
| scr_R | GGCATAGCGACTAAACACATCAAAATACATCTGTGGCTTCACTATTTTGATGTGTTTAGTCGCTA |
| NGS-Lib-Fwd-1 | AATGATACGGCGACCACCGAGATCTA CACTCTTTCCCTACACGACGCTCTTCC GATCTTAAGTAGAGGCTTTATATATCT TGTGGAAAGGACGAAACACC |
| NGS-Lib-Fwd-2 | AATGATACGGCGACCACCGAGATCTA CACTCTTTCCCTACACGACGCTCTTCC GATCTATCATGCTTAGCTTTATATATC TTGTGGAAAGGACGAAACACC |
| NGS-Lib-Fwd-3 | AATGATACGGCGACCACCGAGATCTA CACTCTTTCCCTACACGACGCTCTTCC GATCTGATGCACATCTGCTTTATATAT CTTGTGGAAAGGACGAAACACC |
| NGS-Lib-Fwd-4 | AATGATACGGCGACCACCGAGATCTA CACTCTTTCCCTACACGACGCTCTTCC GATCTCGATTGCTCGACGCTTTATATA TCTTGTGGAAAGGACGAAACACC |
| NGS-Lib-Fwd-5 | AATGATACGGCGACCACCGAGATCTA CACTCTTTCCCTACACGACGCTCTTCC GATCTTCGATAGCAATTCGCTTTATAT ATCTTGTGGAAAGGACGAAACACC |
| NGS-Lib-Fwd-6 | AATGATACGGCGACCACCGAGATCTA CACTCTTTCCCTACACGACGCTCTTCC GATCTATCGATAGTTGCTTGCTTTATA TATCTTGTGGAAAGGACGAAACACC |

| | |
|---|---|
| NGS-Lib-Fwd-7 | AATGATACGGCGACCACCGAGATCTA CACTCTTTCCCTACACGACGCTCTTCC GATCTGATCGATCCAGTTAGGCTTTAT ATATCTTGTGGAAAGGACGAAACACC |
| NGS-Lib-Fwd-8 | AATGATACGGCGACCACCGAGATCTA CACTCTTTCCCTACACGACGCTCTTCC GATCTCGATCGATTTGAGCCTGCTTTA TATATCTTGTGGAAAGGACGAAACAC C |
| NGS-Lib-Fwd-9 | AATGATACGGCGACCACCGAGATCTA CACTCTTTCCCTACACGACGCTCTTCC GATCTACGATCGATACACGATCGCTTT ATATATCTTGTGGAAAGGACGAAACA CC |
| NGS-Lib-Fwd-10 | AATGATACGGCGACCACCGAGATCTA CACTCTTTCCCTACACGACGCTCTTCC GATCTTACGATCGATGGTCCAGAGCTT TATATATCTTGTGGAAAGGACGAAAC ACC |
| NGS-Lib-KO-Rev-1 | CAAGCAGAAGACGGCATACGAGATTCGCCTTGGTGACTGGAGTTCAG ACGTG TGCTCTTCCGATCTCCGACTCGGTGCC ACTTTTTCAA |
| NGS-Lib-KO-Rev-2 | CAAGCAGAAGACGGCATACGAGATATAGCGTCGTGACTGGAGTTCAG ACGTG TGCTCTTCCGATCTCCGACTCGGTGCC ACTTTTTCAA |
| NGS-Lib-KO-Rev-3 | CAAGCAGAAGACGGCATACGAGATGAAGAAGTGTGACTGGAGTTCAG ACGTG TGCTCTTCCGATCTCCGACTCGGTGCC ACTTTTTCAA |
| NGS-Lib-KO-Rev-4 | CAAGCAGAAGACGGCATACGAGATATTCTAGGGTGACTGGAGTTCAG ACGTG TGCTCTTCCGATCTCCGACTCGGTGCC ACTTTTTCAA |
| NGS-Lib-KO-Rev-5 | CAAGCAGAAGACGGCATACGAGATCG TTACCAGTGACTGGAGTTCAGACGTG TGCTCTTCCGATCTCCGACTCGGTGCC ACTTTTTCAA |
| NGS-Lib-KO-Rev-6 | CAAGCAGAAGACGGCATACGAGATGTCTGATGGTGACTGGAGTTCAG ACGTG TGCTCTTCCGATCTCCGACTCGGTGCC ACTTTTTCAA |
| NGS-Lib-KO-Rev-7 | CAAGCAGAAGACGGCATACGAGATTTACGCACGTGACTGGAGTTCAG ACGTG TGCTCTTCCGATCTCCGACTCGGTGCC ACTTTTTCAA |
| NGS-Lib-KO-Rev-8 | CAAGCAGAAGACGGCATACGAGATTTGAATAGGTGACTGGAGTTCAG ACGTG TGCTCTTCCGATCTCCGACTCGGTGCC ACTTTTTCAA |
| NGS-Lib-KO-Rev-9 | CAAGCAGAAGACGGCATACGAGATCGAGTAATGTGACTGGAGTTCAG ACGTG TGCTCTTCCGATCTCCGACTCGGTGCC ACTTTTTCAA |
| NGS-Lib-KO-Rev-10 | CAAGCAGAAGACGGCATACGAGATAATGAGCGGTGACTGGAGTTCAG ACGTG TGCTCTTCCGATCTCCGACTCGGTGCC ACTTTTTCAA |
| GeCKO-F1 | AATGGACTATCATATGCTTACCGTAACTTGAAAGTATTTCG |
| GeCKO-R1 | CTTTAGTTTGTATGTCTGTTGCTATTATGTCTACTATTCTTTCC |

### Supplemental Table 2 – Antibodies

| *Target* | *Company* | *Clone #* | *Catalog #* | *RRID #* |
|---|---|---|---|---|
| pAKT(T308) | Cell Signaling | D25E6 | 13038s | AB_2629447 |
| pAKT(S473) | Cell Signaling | D9E | 4060s | AB_2315049 |
| pAKT(T450) | Cell Signaling | polyclonal | 9267s | AB_823676 |
| AKT | Cell Signaling | polyclonal | 9272s | AB_329827 |
| NICD (V1744) | Cell Signaling | D3B8 | 4147s | AB_2153348 |

| | | | | |
|---|---|---|---|---|
| c-MYC | Abcam | Y69 | ab32072 | AB_731658 |
| P85 | Abcam | EPR18702 | ab191606 | AB_2891324 |
| pS6K (T421/S424) | Cell Signaling | polyclonal | 9204s | AB_2265913 |
| pS6K (T389) | Cell Signaling | polyclonal | 9205s | AB_330944 |
| p70 S6K | Cell Signaling | polyclonal | 9202s | AB_331676 |
| pS6 Ribosomal protein (S235/236) | Cell Signaling | polyclonal | 2211s | AB_331679 |
| S6 Ribosomal protein | Cell Signaling | 54D2 | 2317s | AB_2238583 |
| CRISPR-CAS9 | Abcam | 7A9-3A3 | ab191468 | AB_2692325 |
| BCL-xL | Abcam | polyclonal | ab98143 | AB_10674728 |
| BCL2 | Cell Signaling | polyclonal | 2872T | AB_10693462 |
| TATA binding proteinTBP | Abcam | mAbcam51841 | ab51841 | AB_945758 |
| p-Tuberin/TSC2 (S939) | Cell Signaling | polyclonal | 3615T | AB_2207796 |
| Tuberin/TSC2 | Cell Signaling | D93F12 | 4308T | AB_10547134 |
| pBAD (S136) | Cell Signaling | D25H8 | 4366T | AB_10547878 |
| BAD | Cell Signaling | D24A9 | 9239T | AB_2062127 |
| pRB (S807/811) | Cell Signaling | D20B12 | 8516S | AB_11178658 |
| pRB (S780) | Cell Signaling | polyclonal | 9307T | AB_330015 |
| pRB (S795) | Cell Signaling | unknown | 9301T | AB_330013 |
| RB | Abcam | EPR17512 | ab181616 | AB_2848193 |
| ACTIN | Abcam | 8226 | ab8226 | AB_306371 |
| anti-mouse IgG, HPR-linked | Cell Signaling | polyclonal | 7076S | AB_330924 |
| anti-rabbit IgG, HPR-linked | Cell Signaling | polyclonal | 7074S | AB_2099233 |

### Supplemental Table 3 – Drugs

| Name | Target | CAS | Provider | Catalog # | Delivery |
|---|---|---|---|---|---|
| Palbociclib | CDK4/6:CCND3 | 827022-32-2 | Chemietek | CT-PD2991 | Oral Gavage |
| Ruxolitinib | JAK1/2 | 941678-49-5 | Selleck Chemicals | S1378 | *in vitro* |
| Venetoclax | BCL2 | 1257044-40-8 | LuBioScience | HY-15531 | Oral Gavage |
| MK-2206 | AKT | 1032349-77-1 | LuBioScience | HY-10358 | Oral Gavage |
| Idelalisib | PI3Kdelta | 870281-82-6 | LuBioScience | S2226 | *in vitro* |
| CB-103 | Notch | 218457-67-1 | Chemie Brunschwig AG | CBLSS-5069 | intraperotoneal |
| LY3039478 | Notch | 1421438-81-4 | SpiroChem | S7169 | intraperotoneal |
| DAPT | Notch | 208255-80-5 | Selleck Chemicals | S2215 | *in vitro* |

Supplemental Figure 1. Genome-wide CRISPR screening reveals key candidate pathways synergistic with Notch inhibition.

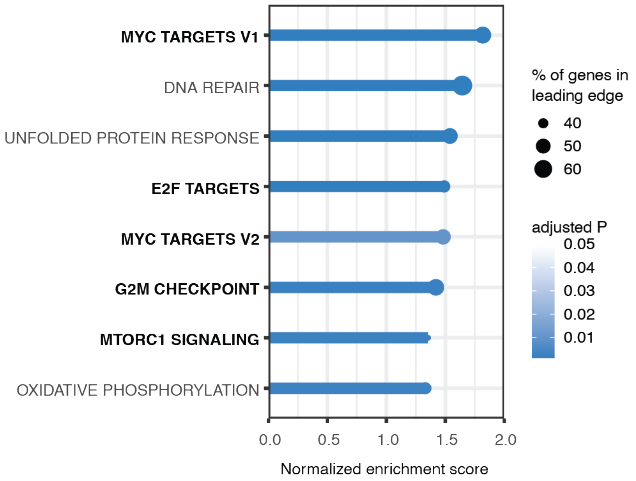


**Supplemental Figure 1. Genome-wide CRISPR screen reveals key candidate pathways synergistic with Notch inhibition.** Gene set enrichment analysis (GSEA) for negative-selection genes both in CB-103 and γ-secretase inhibitor (GSI) treatment in the CRISPR screen based on gene ranking (Robust rank aggregation score) in descending order. Significantly enriched (adjusted $P < 0.05$) hallmark pathways (y-axis) are shown as normalized enrichment score (x-axis). The solid line color scale resembles adjusted *P*, dot size of leadingedge displays percentage of genes enriched in corresponding pathways.

Supplemental Figure 2. Total protein analyses in stable *PIK3R1* knockout or knockdown T-ALL cells.

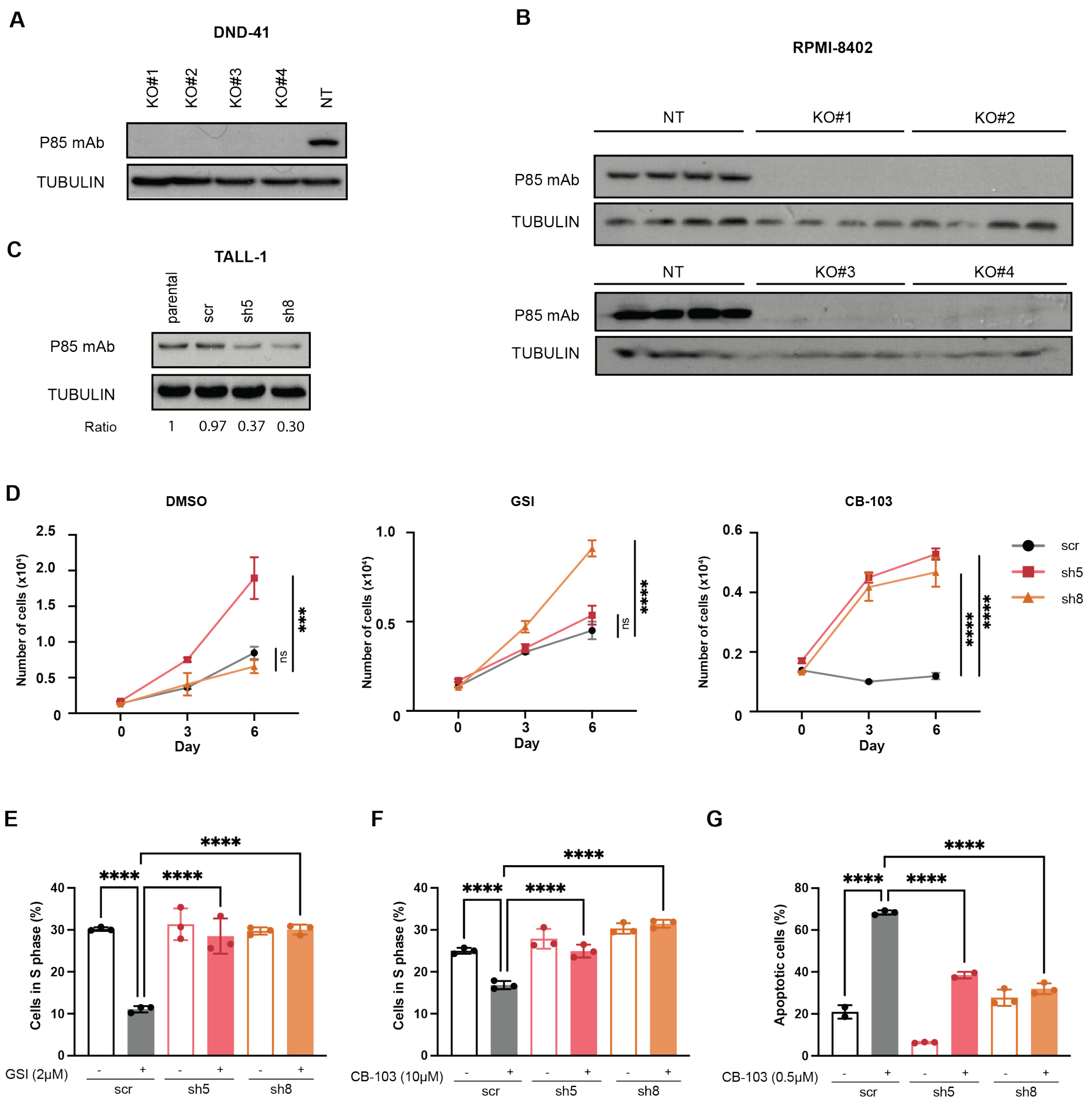


**Supplemental Figure 2. Total protein analyses in stable *PIK3R1* knockout or knockdown T-ALL cells.** (A) Total protein analyses of p85 expression in four different DND-41 *PIK3R1* knockout clones as well as Non-Target (NT) clone is shown. (B) Total protein analyses of p85 expression in four different RPMI-8402 *PIK3R1* knockout clones as well as NT clone is depicted. (C) Total protein analyses of p85 expression in two different TALL-1 *PIK3R1* knockdown cell lines as well as scrambled-hairpin control (scr) line was assessed. Tubublin was used as loading control. (D) Cell proliferation assays of TALL-1 *PIK3R1* knockdown cell lines under DMSO, CB-103 or GSI treatment conditions is presented. Black connected dots, scr line; colored dots (red square – sh5; orange triangle – sh8), *PIK3R1* knockdown cell lines. (E) Cell cycle analyses assessed 6 days post DMSO or GSI treatment at indicated concentration in TALL-1 *PIK3R1* knockdown cell lines (F) Cell cycle analyses assessed 24hrs post DMSO or CB-103 treatment at indicated concentration in TALL-1 *PIK3R1* knockdown cell lines. (G) Apoptosis assay performed 3 days post DMSO or CB-103 treatment at indicated concentration in TALL-1 *PIK3R1* knockdown cell lines. Values shown are mean ± SD (n=3 biologically independent samples, two independent experiments). One-way ANOVA, non-significant (ns), *$P$ value < .0332, **$P$ value < 0.0021, ***$P$ value < 0.0002, ****$P$ value < 0.0001.

Supplemental Figure 3. Experimental design and overview results of RNA-seq study.

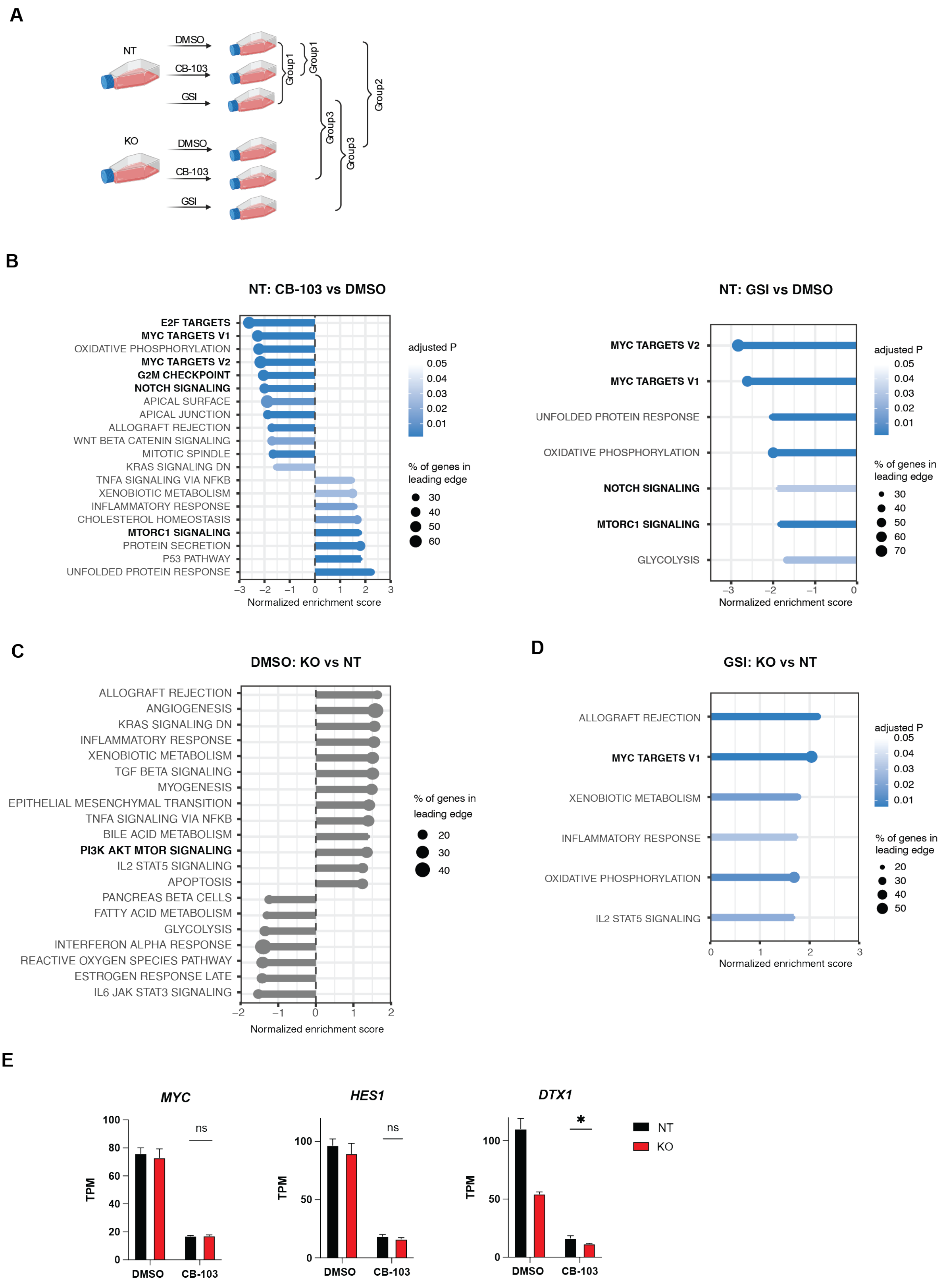


**Supplemental Figure 3. Experimental design and overview results of RNA-seq study.** (A) Experimental set-up and three comparison groups to assess the effect of loss of *PIK3R1* to Notch inhibition in RPMI-8402 cell line at transcriptional level is depicted. (B) Top significantly enriched Hallmark pathways of gene expression level changes comparing CB-103 vs DMSO treated (left panel), GSI vs DMSO treated (right panel) T-ALL cells. (C) Enriched Hallmark pathways of gene expression level changes comparing *PIK3R1* KO vs NT cells. (D) Top significantly enriched Hallmark pathways of gene expression level changes comparing *PIK3R1* KO GSI treated vs NT GSI treated T-ALL cells. Significantly enriched (adjusted $P < 0.05$) hallmark pathways (y-axis) are shown as normalized enrichment score (x-axis). The solid line color scale resembles adjusted $P$ (grey, non-significant), dot size of leadingedge displays percentage of genes enriched in corresponding pathways. (E) Expression of indicated genes measured as transcripts per million (TPM) comparing *PIK3R1* KO (red bars) vs NT (black bars) cells with or without CB-103 treatment. Values shown are mean $\pm$ SD. One way ANOVA test, *$P$ value < .0332, **$P$ value < 0.0021, ***$P$ value < 0.0002, ****$P$ value < 0.0001.

Supplemental Figure 4. Experimental design and overview of proteomics and phosphoproteomics study.

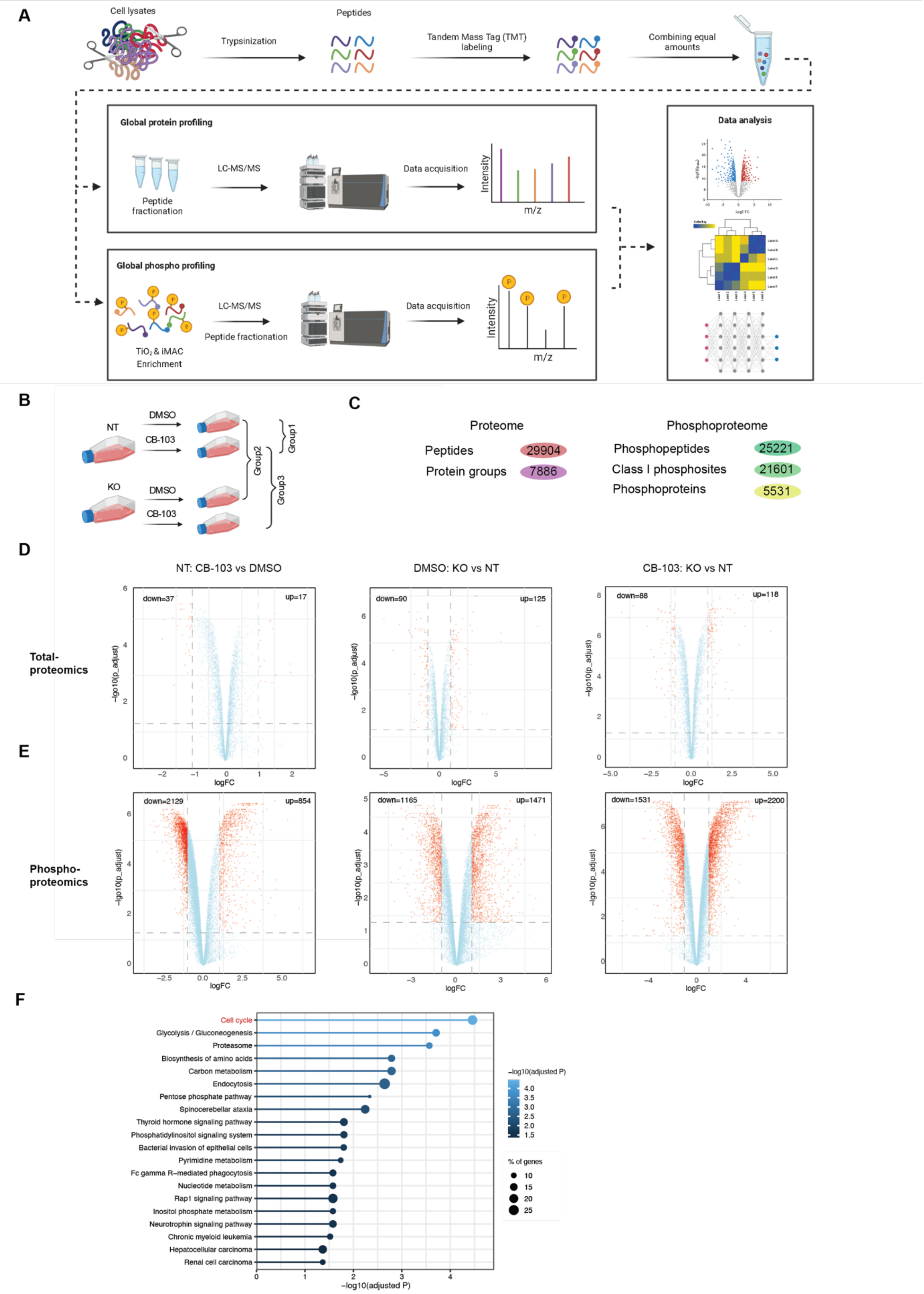


**Supplemental Figure 4. Experimental design and overview of total proteomics and phosphoproteomics study.** (A) Workflow of total proteomics and phosphoproteomics study. (B) Experimental set-up and three comparison groups to assess the effect of loss of *PIK3R1* to

CB-103 in RPMI-8402 T-ALL cell line. (C) Overview of results from proteome and phosphoproteome analysis. (D) Volcano plot depicting protein level changes for each comparison. Red, significant, FDR < 0.05, abs ($\log_2$ fold-change)>1; blue, non-significant (ns). (E) Volcano plot displaying phosphorylational level changes for each comparison. Red, significant, FDR < 0.05, abs ($\log_2$ fold-change)>1; blue, non-signficant (ns). (F) Significantly enriched KEGG pathways of proteins with altered total protein level changes comparing *PIK3R1* KO CB-103 treated vs NT CB-103 treated T-ALL cells. Top 20 pathways are shown, the solid line color scale resembles adjusted *P*, dot size of leading edge displays percentage of genes enriched in corresponding pathways.

Supplemental Figure 5. DND-41 *PIK3R1* KO cells obtain altered phosphorylation profiles in response to Notch inhibition.

A

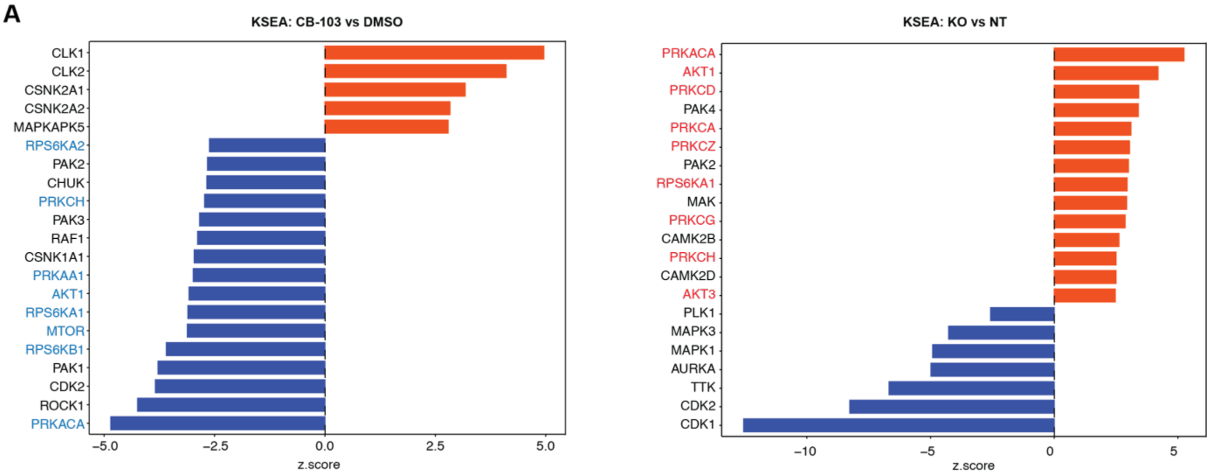


B

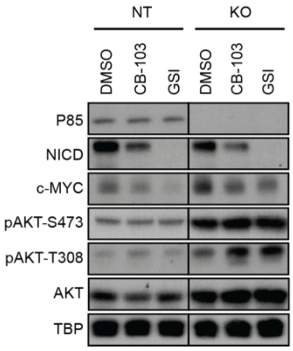


**Supplemental Figure 5. DND-41 *PIK3R1* KO cells obtain altered phosphorylation profiles in response to Notch inhibition.** (A) Kinase-Substrate-Enrichment-Analysis (KSEA) of phosphorylation profiles comparing CB-103 vs DMSO (left panel) and *PIK3R1* KO vs NT (right panel) RPMI-8402 cells. Red, kinases with positive z-score; blue, kinases with negative z-score. (B) Total protein and phosphorylation level of indicated phosphosites by Western blotting for key proteins involved in Notch and PI3K-AKT pathways in DND-41 NT or PIK3R1 KO cells. TATA-box binding protein (TBP) was used as loading control.

Supplemental Figure 6. Phosphoproteomics analysis of *PIK3R1* KO cells reveals altered phosphorylation profiles of splicing factors in response to Notch inhibition.

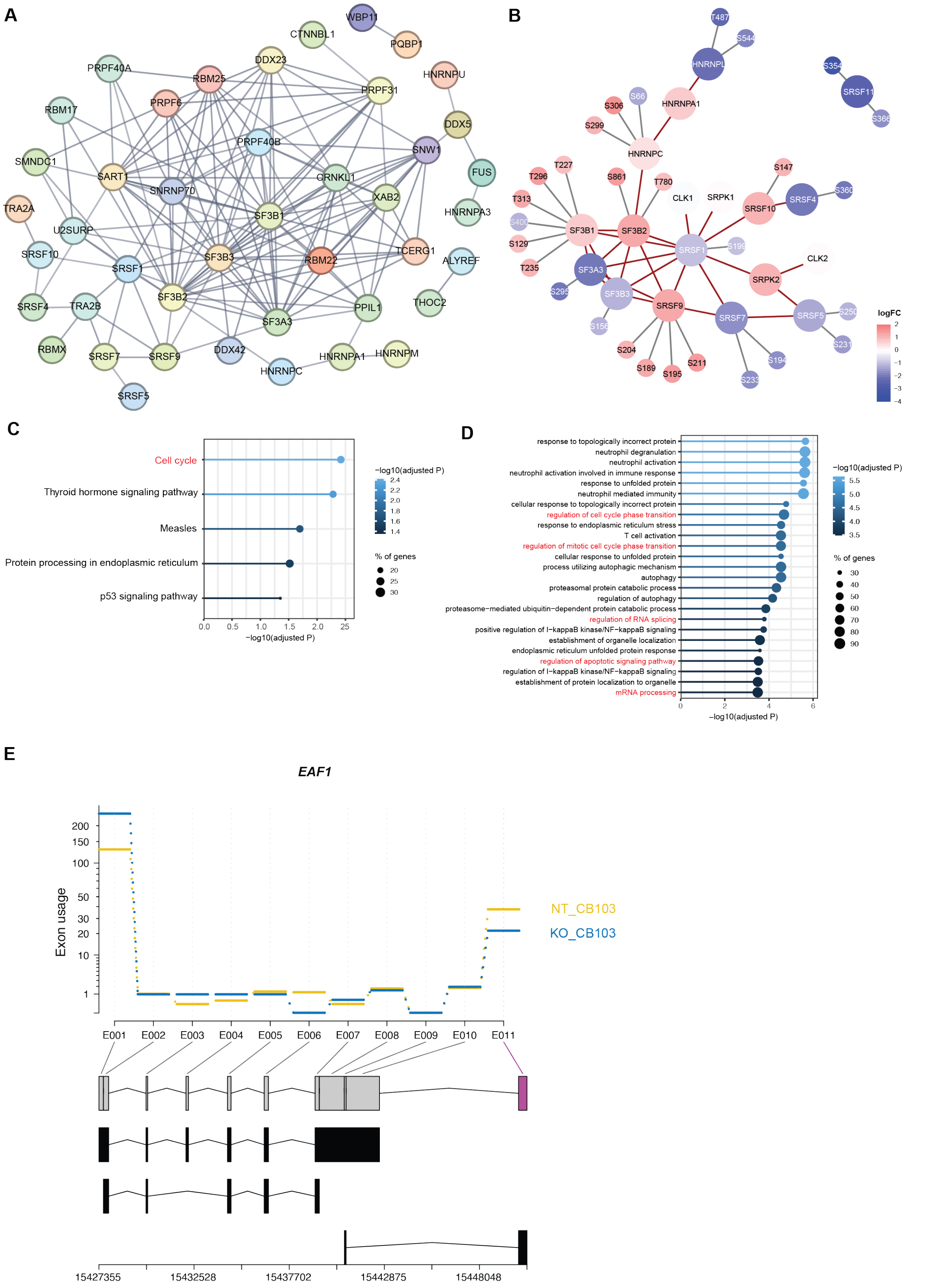


**Supplemental Figure 6. Phosphoproteomics analysis of *PIK3R1* KO cells reveals altered phosphorylation profiles of splicing factors in response to Notch inhibition.** (A) STRING analysis of phosphoproteins within Splicesome pathway in Figure 4A. Line thickness indicates the strength of data support ("Confidence"), interaction source is "Experiments", minimum required interaction score is 0.700. (B) Detailed plots of key phosphoproteins with annotated phosphosites and corresponding fold changes from A. Red circles identified phosphoproteins with phosphorylational changes as fold-change > 2, FDR < 0.05. Grey circles identified phosphoproteins with phosphorylation sites omitted. Red connecting line, interaction with proteins demonstrated by String analyses, grey radiating line, depicting detailed phosphorylation sites associated with phosphoproteins. (C) Top significantly enriched KEGG pathways of transcripts with altered expression levels comparing *PIK3R1* KO CB-103 treated vs NT CB-103 treated T-ALL cells. The solid line color scale resembles adjusted *P*, dot size of leadingedge displays percentage of genes enriched in corresponding pathways. (D) Top significantly enriched gene ontology biological process (GOBP) pathways of transcripts with altered expression levels comparing *PIK3R1* KO CB-103 treated vs NT CB-103 treated T-ALL cells. The solid line color scale resembles adjusted *P*, dot size of leadingedge displays percentage of genes enriched in corresponding pathways. (E) Fitted splicing plot from DEXSeq analysis displaying differential exon usages of gene *ELL Associated Factor 1* (*EAF1*) comparing *PIK3R1* KO CB-103 treated vs NT CB-103 treated T-ALL cells. Purple, altered exon usage.
